# A unique epigenomic landscape defines CD8^+^ tissue-resident memory T cells

**DOI:** 10.1101/2022.05.04.490680

**Authors:** Frank A. Buquicchio, Raissa Fonseca, Julia A. Belk, Maximilien Evrard, Andreas Obers, Yanyan Qi, Bence Daniel, Kathryn E. Yost, Ansuman T. Satpathy, Laura K. Mackay

## Abstract

Memory T cells provide rapid and long-term protection against infection and tumors. The memory CD8^+^ T cell repertoire contains phenotypically and transcriptionally heterogeneous subsets with specialized functions and recirculation patterns. While these T cell populations have been well characterized in terms of differentiation potential and function, the epigenetic changes underlying memory T cell fate determination and tissue-residency remain largely unexplored. Here, we examined the single-cell chromatin landscape of CD8^+^ T cells over the course of acute viral infection. We reveal an early bifurcation of memory precursors displaying distinct chromatin accessibility and define epigenetic trajectories that lead to a circulating (T_CIRC_) or tissue-resident memory T (T_RM_) cell fate. While T_RM_ cells displayed a conserved epigenetic signature across organs, we demonstrate that these cells exhibit tissue-specific signatures and identify transcription factors that regulate T_RM_ cell populations in a site-specific manner. Moreover, we demonstrate that T_RM_ cells and exhausted T (T_EX_) cells are distinct epigenetic lineages that are distinguishable early in their differentiation. Together, these findings show that T_RM_ cell development is accompanied by dynamic alterations in chromatin accessibility that direct a unique transcriptional program resulting in a tissue-adapted and functionally distinct T cell state.

**Graphical Abstract:** 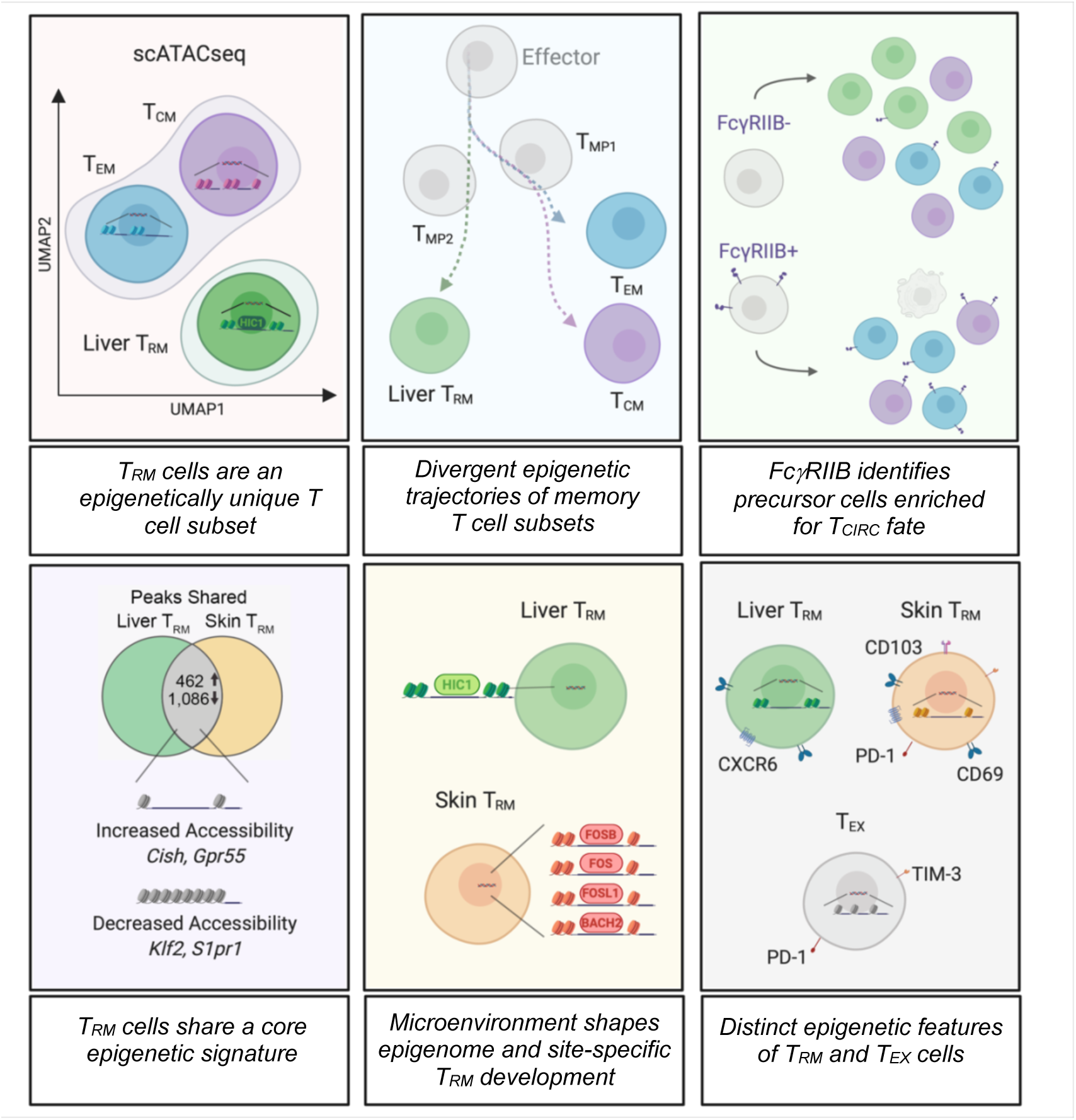

**Highlights:**

- scATAC atlas reveals the epigenetic variance of memory CD8^+^ T cell subsets over the course of acute infection
- Early bifurcation of memory precursors leads to circulating versus tissue-resident cell fates
- Integrating transcriptional and epigenetic analyses identified organ-specific T_RM_ cell regulators including HIC1 and BACH2
- Epigenetic distinction of T_RM_ cells and T_EX_ cell subsets

## Introduction

CD8^+^ T cells are key mediators of protective immunity against infectious diseases and tumors. Following antigen encounter, activated T cells can infiltrate sites of infection where they mediate pathogen control. Disease resolution is followed by the generation of heterogeneous memory T cells with specialized functions and recirculation patterns. Whereas some memory T cells circulate throughout the blood and lymphatics (including central (T_CM_) and effector (T_EM_) memory T cells), others are permanently stationed in peripheral organs. These non-migratory tissue-resident memory T (T_RM_) cells have been identified in virtually all tissues across species, and are critical for infection and cancer control (Masopust and Soerens, 2019; Okla et al., 2021).

Over recent years, high-throughput sequencing technologies have been used to define transcriptional and epigenetic changes associated with T cell differentiation in various settings (Joshi et al., 2007; Kallies et al., 2009; Mackay et al., 2016; Milner et al., 2017, 2020; Pauken et al., 2016; Roychoudhuri et al., 2016; Sen et al., 2016; Skon et al., 2013; Utzschneider et al., 2020; Zhou et al., 2010). Gene expression analyses and protein profiling have demonstrated that variations in tissue microenvironments, cytokine exposure, antigen persistence, and TCR signal strength poise T cells at different stages and shape the effector response, culminating in the development of heterogeneous circulating memory (T_CIRC_) and T_RM_ cell subsets (Beura et al., 2018; Christo et al., 2021; Joshi et al., 2007; Kumar et al., 2017; Mackay et al., 2016; Masopust et al., 2010; Sarkar et al., 2008; Solouki et al., 2020; Wakim et al., 2010). These populations of memory T cells display distinct stemness and functional abilities, and together provide optimal immune protection and long-term immunity against pathogens and tumors (Behr et al., 2020; Christo et al., 2021; Fonseca et al., 2020; Jameson and Masopust, 2009; Kaech and Cui, 2012; Park et al., 2019; Sallusto et al., 1999). In certain contexts, such as chronic infections or persistent tumors, continuous T cell stimulation prompts terminal differentiation leading to the development of exhausted T (T_EX_) cells that display increased expression of inhibitory receptors and reduced functional ability as compared to memory T cell populations (Im et al., 2016; Mackay et al., 2012a; McLane et al., 2019; Miller et al., 2019; Paley et al., 2012).

Accordingly, dynamic genome-wide changes in DNA methylation and chromatin accessibility observed during T_CIRC_ development has demonstrated the complex determination of effector and memory T cell fates in both mice and humans (Akondy et al., 2017; Araki et al., 2009; Scharer et al., 2013). Most recently, epigenomic profiling of single cells has demonstrated changes in *cis*- and *trans*-regulatory elements associated with regulation of gene expression in individual cell types, allowing for reconstruction of trajectories associated with cellular differentiation (Lareau et al., 2019; Satpathy et al., 2019). While it is known that heterogeneous T_CIRC_, T_EX_ and T_RM_ cells exist and rely on distinct transcriptional circuitries, the epigenomic changes steering their ontogeny, as well as when their differentiation trajectories diverge is not known. Moreover, our understanding of the collective epigenetic variation of T_RM_ cells across different tissues remains unexplored.

Here, we used the single-cell assay for transposase accessible chromatin with sequencing (scATAC-seq) to examine the dynamic genome-scale changes in chromatin accessibility that occur in CD8^+^ T cells over the course of viral infection. We found subset-specific variations in the epigenetic landscape of memory T cells that are generated in response to acute LCMV infection, with differences in *cis*-regulatory element accessibility being established early post-infection in memory precursor populations. Increased *Fcgr2b* locus accessibility reflected the dynamics of Fc*γ*RIIB expression in T cells *in vivo*, allowing the selection of memory precursors with enhanced capacity to generate either T_CIRC_ or T_RM_ cells in the liver. Whereas T_RM_ cells were epigenetically distinct from their circulating counterparts and exhibited conserved features between tissues, our analyses also revealed transcription factors that regulate T_RM_ cell formation in a tissue-specific manner. Moreover, despite confirming considerable phenotypic similarities between T_RM_ and T_EX_ cells, we demonstrated that T_RM_ cells display a distinct chromatin landscape that share relatively few features with T_EX_ cell subsets. Together, our data indicate that distinct epigenetic landscapes accompany memory T cell differentiation and form the basis of the transcriptional and functional differences associated with unique T cell ontogenies. Additionally, we provide a <u>public genome browser</u> for interrogating the chromatin accessibility profiles of effector and memory CD8^+^ T cell populations, which will facilitate future investigations.

## Results

### T_RM_ cells are an epigenetically distinct subset of memory T cells

To investigate the epigenetic landscape of CD8^+^ memory T cell populations, we first utilized a model of acute LCMV infection (Armstrong strain) in combination with adoptive transfer of TCR transgenic CD8^+^ T cells. To this end, we transferred naïve congenically labelled CD45.1^+^ P14 cells specific for LCMVgp33-41 into C57BL/6 mice that were infected with LCMV to generate antigen-specific memory CD8^+^ T cells across organs. At 30 d post infection (p.i.), T_EM_ and T_CM_ P14 cells were isolated from the spleen, and T_EM_, T_CM_ and T_RM_ P14 cells were isolated from the liver, alongside P14 cells from naive animals (**Figure S1A**). Naïve and memory P14 cell populations were then subjected to scATAC-seq (Satpathy et al., 2019) (**Figure 1A**). In total, 15,740 single cells passed quality control filters of at least 1,000 unique fragments per cell and a transcription start site (TSS) enrichment score greater than or equal to 5 (**Figure S1B**). To analyse memory T cell epigenetic profiles, we utilised ArchR (Granja et al., 2021): (1) for dimensionality reduction using Latent Sematic Indexing (LSI), Uniform Manifold Approximation and Projection (UMAP) embedding and cell clustering(Becht et al., 2019; Cusanovich et al., 2015); (2) to analyze changes in accessibility of individual regions in the genome (‘peaks’); (3) to calculate deviation in accessibility for transcription factor (TF) motifs; (4) to visualize aligned ATAC-seq reads; (5) to predict ‘marker peaks/genes’ or changes in accessibility that are specific to a given cluster; (6) to model differential gene expression using gene activity score (‘gene score’) and (7) to perform pseudo-time differentiation trajectory analysis to model epigenetic changes that occur over the course of a projected differentiation trajectory (**Figure 1B**).

**Figure 1.**
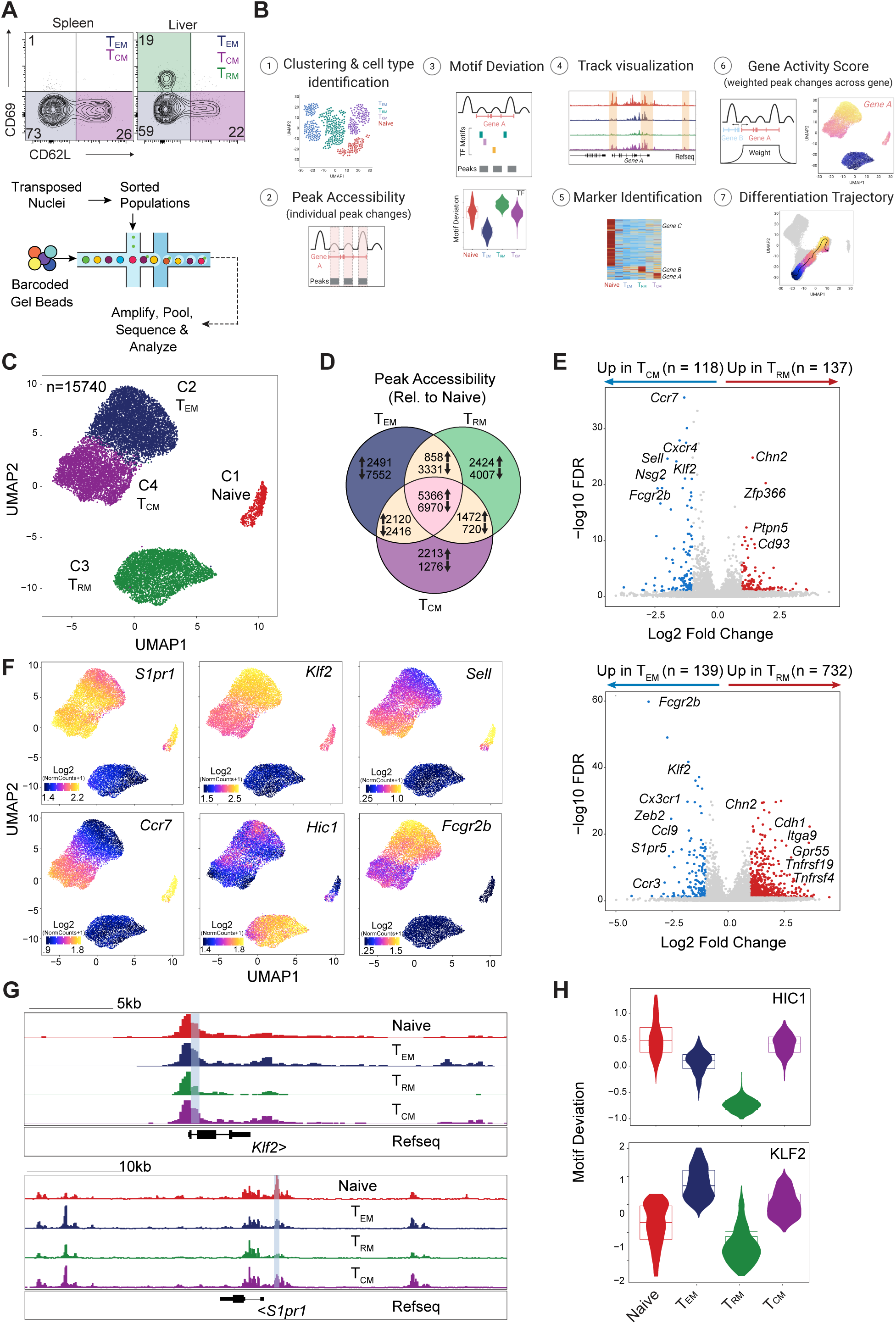
T_RM_ cells display a unique epigenetic landscape amidst memory T cell subsets. **(A-H)** Congenically marked naïve CD8^+^ P14 cells were transferred into C57Bl/6 naïve recipient mice followed by LCMV Armstrong infection. T_CM_ (CD62L^+^ CD69^-^), T_EM_ (CD62L^-^CD69^-^) and T_RM_ (CD62L^-^CD69^+^) cells were flow sorted from the spleen and liver 30 d p.i. and scATACseq was performed. **(A)** Experimental schematics of scATACseq droplets and representative flow plots of sorted populations from spleen and liver. **(B)** Analyses performed in single-cell chromatin accessibility data using ArchR. **(C)** UMAP projection of memory T cells. **(D)** Venn diagram of differential peaks in identified clusters individually compared to naive cluster (log2 FC > 1, FDR > 10). **(E)** Gene score volcano plots identifying genes with significantly different accessibility (log2 FC > 1, FDR > 10) between T_RM_ and T_EM_ or T_CM_ memory clusters; notable genes annotated manually. **(F)** UMAP depicting relative gene accessibility (gene score) across clusters. **(G)** *Klf2* and *S1pr1* genome tracks (height normalized) and **(H)** KLF2 and HIC1 motif deviation in indicated clusters.

T cell visualization by UMAP revealed four major clusters (**Figure 1C**). These projected clusters nearly exclusively aligned with sorted T cell subsets, suggesting that scATAC-seq accurately identifies the epigenetic heterogeneity within CD8^+^ memory T cells (**Figure S1C**). Thus, cluster identity was assigned according to the sorted T cell population represented in each cluster (naïve T cells (C1), T_EM_ (C2), T_RM_ (C3) and T_CM_ (C4) cells). Whereas our analyses revealed comparatively little distinction between the chromatin state of T_CM_ and T_EM_ cells regardless of tissue origin, T_RM_ cells derived from the liver comprised a discrete cluster, positioned separately from the T_CIRC_ cell populations (**Figure 1C**). To identify epigenetic differences between T_CIRC_ and T_RM_ cells, we compared each memory T cell cluster to naïve T cells to identify differences in peak accessibility and compared these peak sets across the three memory clusters. This analysis showed that each cluster exhibited unique peak accessibility changes that were not observed in the other respective clusters, as well as a peak set that was shared across memory T cells. Importantly, the number of peaks exclusive to the T_RM_ cell cluster (2,424 increased, 4,007 decreased) was similar in magnitude to the number of T_CM_ (2,213, 1,276) and T_EM_ (2,491, 7,552) cell-exclusive peaks, further confirming that T_RM_ cells are an epigenetically distinct subset of memory T cells, while T_CM_ and T_EM_ cells shared proportionally higher number of peaks (2,120, 2,416), supporting the increased epigenetic similarity observed between these clusters (**Figure 1D**).

To link differentially accessible ATAC-seq peaks to genes, we examined gene-level accessibility changes, both across the gene body and in linked distal sites, by gene score(Granja et al., 2021; Pliner et al., 2018). Direct comparison of gene scores between T_RM_ and T_CM_ and/or T_EM_ cell clusters revealed the expected differential accessibility in peaks corresponding to *Klf2, Ccr7, Sell, S1pr5, Zeb2* and *Cx3cr1*, genes all known to be downregulated in T_RM_ cells as compared to circulating populations (**Figure 1E**). Importantly, the downregulation of KLF2 and its target S1PR1, as well as ZEB2 and S1PR5, are known to be critical to halt tissue egress in order for T_RM_ cells to develop in organs (Evrard et al., 2022; Mackay et al., 2013; Skon et al., 2013). Our data also revealed the differential accessibility of genes with no known role in liver T_RM_ cell formation such as *Fcgr2b* and *Hic1*, as well as increased accessibility of adhesion-related genes such *Chn2*, *Cdh1, Itga9,* and *Gpr55*, a G-protein coupled receptor known to regulate intraepithelial lymphocyte (IEL) migration (Sumida et al., 2017) in T_RM_ cells (**Figure 1E and 1F**). Diminished gene scores of *Ccr7* and *Sell* were also observed in T_EM_ relative to T_CM_ clusters (*Sell*; L2fc = -.327, FDR = 2.45. *Ccr7*; L2fc = -.820, FDR = 24.20) as well as increased accessibility in *S1pr5* and *Zeb2* loci in T_EM_ cells (**Figure S1D and S1E**) in line with the anticipated expression pattern of these molecules.

Reduced accessibility at the *Klf2* and *S1pr1* loci in T_RM_ cells was accompanied by decreased KLF2 motif accessibility (**Figure 1G and 1H**). Conversely, while the *Hic1* locus displayed increased accessibility in T_RM_ cells, the HIC1 motif was significantly less accessible in this population, consistent with its role as a transcriptional repressor (Pinte et al., 2004) (**Figure 1F and 1H**). Similarly, the transcriptional repressor BLIMP-1, encoded by *Prdm1*, displayed decreased motif accessibility in the T_EM_ cell subset and increased *Prdm1* gene score (**Figure S1F-S1H**), supporting previous findings that demonstrated the expression and major role for this TF in T_EM_ development by promoting CD8_+_ T cell proliferative response and differentiation (Rutishauser et al., 2009). Moreover, while reduced expression of BLIMP-1 is observed in T_RM_ cells, BLIMP-1 and HOBIT deficiency was shown to be detrimental for LCMV-specific T_RM_ cell formation in the liver (Mackay et al., 2016). Together, our data demonstrate that T_RM_ cells are an epigenetically distinct memory T cell subset and the ability of scATAC-seq to identify unique features of memory T cell subsets at the chromatin level.

### Memory T cell subsets display distinct epigenetic trajectories

Recent evidence suggests that T_RM_ cell fate is determined early during infection (Kok et al., 2020, 2021; Kurd et al., 2020; Mani et al., 2019; Milner et al., 2017). We sought to understand the progression of epigenetic changes during T cell differentiation and to determine whether progenitors that preferentially give rise to either T_CIRC_ or T_RM_ cell populations could be identified. For this, we focused our analyses on the liver, an organ comprising T_RM_, T_EM_ and T_CM_ cell population (Fernandez-Ruiz et al., 2016), and sorted P14 cells at 7, 14, and 30 d following LCMV infection, alongside T_EM_ and T_CM_ P14 cells from the spleen. At 14 d p.i., we sorted both CD69^-^ and CD69^+^ cells from the liver to more finely define the CD69^+^ T_RM_ cell-poised population (**Figure S2A**). As above, samples were subjected to scATAC-seq, analyzed, and visualized by UMAP alongside naïve P14 cells from non-infected mice (**Figure 2A**).

**Figure 2.**
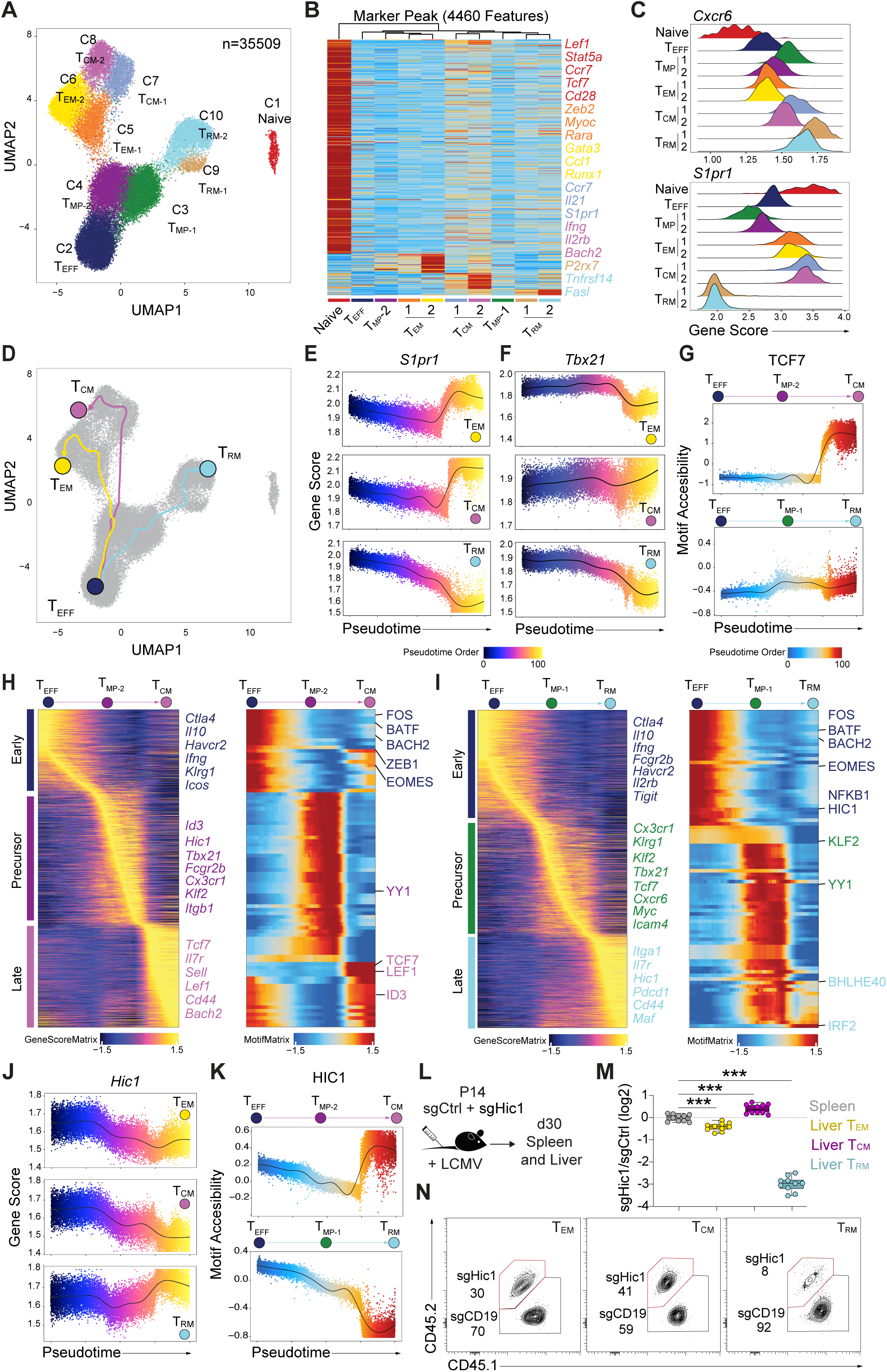
Distinct epigenetic trajectories define T_CIRC_ and T_RM_ cell development. **(A-K)** Congenically marked naïve CD8^+^ P14 cells were transferred into C57Bl/6 naïve recipient mice followed by LCMV Armstrong infection. P14 cells were sorted from the liver at 7 d (total P14 cells), 14 d (CD69^-^ and CD69^+^), alongside T_RM_ (CD62L^-^CD69^+^), T_CM_ (CD62L_+_ CD69^-^) and T_EM_ (CD62L^-^CD69^-^) from the liver and spleen at 30 d p.i. and scATACseq was performed. **(A)** UMAP projection of scATAC profiles of flow sorted populations. **(B)** Marker peak heatmap identifying cis-regulatory elements uniquely active in individual clusters; peaks in or linked to notable genes are annotated and colored by cluster. **(C)** *Cxcr6* and *S1pr1* gene scores in individual clusters. **(D)** Predicted differentiation trajectories of identified memory clusters. **(E)** *S1pr1* and **(F)** *Tbx21* gene score over pseudotime for individual trajectories. **(G)** Motif accessibility of TCF7 over pseudotime during T_CM_ and T_RM_ cell epigenetic trajectories. **(H)** Heatmaps with dynamic gene score (left) and **(I)** motif accessibility (right) over pseudotime during T_CM_ and T_RM_ differentiation trajectories; notable genes and motifs that appear in the trajectory are annotated near their approximate position in pseudotime. **(J)** *Hic1* gene score and **(K)** motif accessibility over pseudotime. **(L-N)** Control (sgCtrl) or *Hic1* (sgHic1) ablation was performed using CRISPR-Cas9 in distinct congenically marked naïve CD8^+^ P14 cells. Cells were then transferred into LCMV infected recipients and isolated from the spleen and liver 30 d p.i. **(L)** Experimental schematics. **(M)** Log2 fold change of sgHic1 and sgCtrl indicated cell subsets normalized to the spleen and **(N)** Representative flow plots of transferred cells for the indicated subsets in liver. Data is representative from **(L-N)** 2 independent experiments with n=10 mice each. In **(M)** symbols represent individual mice. Box plots show the median, interquartile range and minimum/maximum whiskers. *** p≤0.001, One-way ANOVA with Bonferroni post-test.

We found that early effector P14 cells isolated at 7 d p.i. were highly heterogenous and fell into three distinct clusters (C2, C3, C4). Whereas C2 was comprised almost solely of 7 d p.i. cells and denoted as effector T cells (T_EFF_) (by abundance at 7 d p.i. and by loss of accessibility at the *Il7r* and *Il2* loci), other cells isolated from this timepoint clustered together with cells isolated at 14 d p.i. in C3 or C4 (**Figure 2A and S2B**) indicating the early divergence of T_RM_ and T_CIRC_ precursors following infection. We denoted C3 and C4 as memory precursor cells (T_MP_); CD69_+_ cells isolated at 14 d p.i. were nearly exclusively contained within C3 (T_MP-1_), indicating close relation to the T_RM_-exclusive clusters, whereas CD69_-_ samples were distributed between C3 and C4 (T_MP-1_ and T_MP-2_, respectively; **Figure S2B**). We identified genes with cluster-specific accessibility increases in potential *cis*-regulatory elements, including *Zeb2* in T_EM_ cells, *Ccr7 in* T_CM_ cells, and *Fasl* and *P2rx7* in T_RM_ cells (**Figure S2B**). Accordingly, correlation analysis of these marker peaks supported the epigenetic similarity of T_MP-1_ and T_RM_ cells indicating an association between these populations (**Figure 2B**). Moreover, T_MP-1_ exhibited a higher gene score in the T_RM_ cell-associated gene *Cxcr6* (Fernandez-Ruiz et al., 2016; Wein et al., 2019) and decreased gene score in *S1pr1* when compared to T_MP-2_, further suggesting that the T_MP-1_ cluster precedes T_RM_ cell differentiation (**Figure 2C**).

Next, we constructed differentiation trajectories using the T_EFF_ cluster as a starting point to computationally order T cell clusters along a pseudotime axis. Here, single cells are aligned to the trajectory by calculating the nearest cell-to-trajectory distance, which allows cells to be ordered by their determined position in pseudotime (Granja et al., 2021). Our analysis thus far suggested that T_MP-1_ (C3) preceded T_RM_ cell development, whereas T_MP-2_ (C4) bore closer similarity to T_CIRC_ (T_EM_ and T_CM_) cell clusters. As such, our pseudotime trajectories navigated through these T_MP_ clusters, from T_EFF_ *→*T_MP-1_ *→*T_RM_, or from T_EFF_ *→*T_MP-2_ *→*T_CM_ and T_EM_ cell populations (**Figure 2D**).

We observed several expected changes along these developmental trajectories, including a progressive loss in *S1pr1* gene score along the T_EFF_ *→*T_RM_ trajectory, coupled with increased *S1pr1* gene score from T_EFF_ *→*T_CM_ and T_EFF_ *→*T_EM_ (**Figure 2E**). Further, *Cxcr6* and *Sell* followed anticipated trends, with *Cxcr6* gene score increasing in the T_EFF_ *→*T_RM_ trajectory, while remaining relatively consistent in T_EFF_ *→*T_EM_. *Sell* accessibility decreased from T_EFF_ *→*T_RM_ but remained accessible in the T_EFF_ *→*T_CM_ trajectory (**Figure 2C and 2D**). Gene accessibility changes in *Sell* and *Cxcr6* were reflective of protein expression changes observed by flow cytometry at the same time points (**Figure S2E**). Trajectory analysis demonstrated a decrease in the *Tbx21* (encoding T-BET) gene score along the T_EFF_ *→*T_RM_ trajectory (**Figure 2F**), aligning with previous work showing T-BET downregulation during T_RM_ cell development (Laidlaw et al., 2014; Mackay et al., 2015). Additionally, we found an increase in the TCF7 motif accessibility along the T_EFF_ *→* T_CM_ trajectory, corroborating the known role of this TF in promoting T_CM_ differentiation (Gattinoni et al., 2009; Jeannet et al., 2010) (**Figure 2G**).

Global changes in gene score and motif accessibility across pseudotime from T_EFF_ to either T_CM_ or T_RM_ cells revealed several patterns between these developmental pathways. In the T_EFF_ *→*T_MP-2_ *→*T_CM_ trajectory, the accessibility of *Id3, Tcf7*, *Lef1*, and *Sell* progressively increased along this path, together with increased motif accessibility in differentiated T_CM_ as anticipated (**Figure 2H**). In the T_EFF_ *→*T_MP-1_ *→*T_RM_ trajectory, *Cx3cr1* and *Klrg1* accessibility was lost early in the T_EFF_ *→* T_MP-1_ transition, consistent with evidence showing the inability of CX3CR1^+^KLRG1^+^ effector T cells to give rise to T_RM_ cells (Gerlach et al., 2016; Herndler-Brandstetter et al., 2018) (**Figure 2I**). Similarly, KLF2 motif accessibility was lost along the T_RM_ cell transition whereas BHLHE40 motif accessibility was gained during the T_MP-1_ to T_RM_ cell transition (**Figure 2I**), consistent with the observed role of BHLHE40 in this subset (Li et al., 2019). Consistent with changes observed in cells isolated from the liver at day 30 post infection, an increase in *Hic1* gene score was observed in T_RM_ cells over pseudotime (**Figure 2J**), followed by a reduction in motif accessibility of HIC1 sites in T_RM_ cells, with opposite trends in T_EFF_ *→*T_CM_ (**Figure 2K**), suggesting that HIC1 represses target gene accessibility early in T_RM_ cell differentiation.

While HIC1 is known to regulate the IEL population in the small intestine (Burrows et al., 2017), whether this transcription factor is required for liver T_RM_ cell development is not known. To investigate the functional relevance of HIC1 in liver T_RM_ cell development, we used CRISPR-Cas9 to ablate HIC1 in P14 cells that were subsequently transferred into LCMV infected recipients (**Figure 2L**). Importantly, we found that the genetic deletion of HIC1 resulted in significant reduction of CD69^+^ T cells in the liver as early as 9d p.i. (**Figure S2F-S2H**). At 30d p.i., HIC1-deficient liver T_RM_ cells were further depleted, with decreased effect of HIC deletion on T_CIRC_ populations (**Figure 2M-2N and S2I**), suggesting that HIC1 is a critical regulator of T_RM_ cell differentiation. Together, these data support previous findings (Kok et al., 2020; Kurd et al., 2020; Milner et al., 2020) and reinforce the notion that T_RM_ cell fate may sealed early post T cell activation (Kok et al., 2021).

### FcγRIIB expression identifies precursors enriched for the T_CIRC_ cell fate

Our trajectory analyses indicated that a subset of T cells may be poised for T_RM_ cell differentiation at early stages following infection. To identify surface markers that would allow for the isolation of putative T_RM_ cell precursors, we investigated genes encoding cell surface markers with differential accessibility in T_MP-1_ relative to T_MP-2_ cell clusters (**Figure 3A**). To account for potential heterogeneity within the T_MP_ clusters, we also accounted for whether these genes were also differentially accessible in fully differentiated T_RM_ cells relative to T_CIRC_ cells. Among the genes observed, *Cx3cr1* and *S1pr5* showed increased gene scores in T_MP-2_ and T_CIRC_ relative to T_MP-1_ and T_RM_ cells, indicating the presence of an effector population poised for T_CIRC_ cell differentiation (**Figure 3B**). We also observed significantly decreased accessibility of *Fcgr2b* (FcγRIIB) in both T_MP-2_ and T_RM_ cells relative to T_MP-1_ and T_CIRC_ cells (**Figure 3B and S3A**). FcγRIIB is a low-affinity Fc receptor known to act as an inhibitory receptor in CD8^+^ T cells and promote apoptosis(Morris et al., 2020; Starbeck-Miller et al., 2014), although its role in T_RM_ cell differentiation has not been explored.

**Figure 3.**
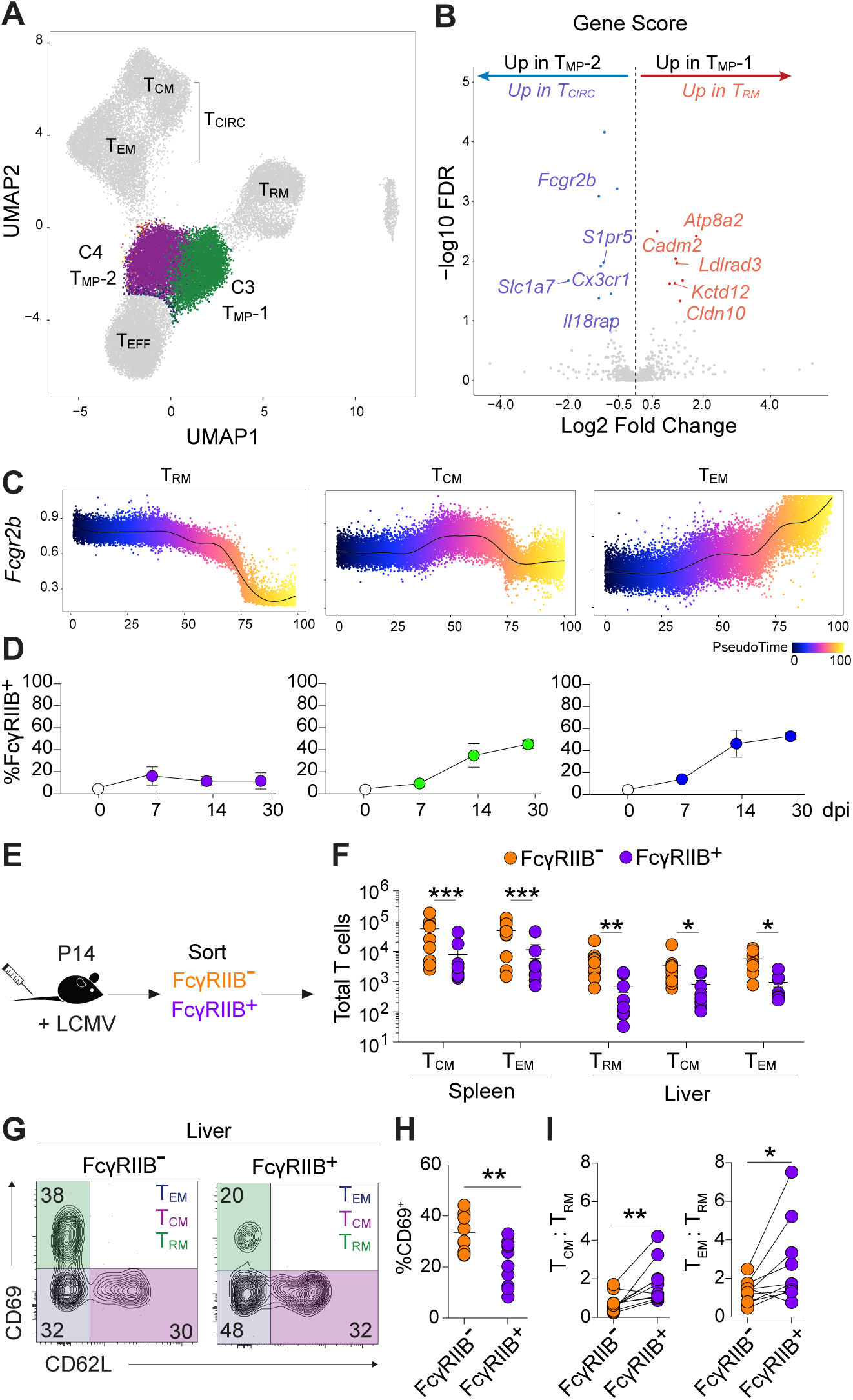
FcγRIIB expression identify memory precursors with T_CIRC_ cell differentiation bias. **(A-C)** Congenically marked naïve CD8^+^ P14 cells were transferred into C57Bl/6 naïve recipient mice followed by LCMV Armstrong infection. P14 cells were flow sorted: from the liver at 7 d p.i., CD69- and CD69^+^ at 14 d p.i. and T_CM_, T_EM_ and T_RM_ from the spleen and liver at 30 d p.i. and scATACseq was performed. **(A)** UMAP projection of scATACseq profiles of flow sorted populations with memory T cell precursor clusters (T_MP_-1 and T_MP_-2) highlighted. **(B)** Gene score volcano plots identifying genes with significantly different accessibility (log2 fold change > 1, FDR > 10) between T_CIRC_ and T_RM_ clusters with similar changes in T_MP_-2 and T_MP_-1 clusters; notable genes were annotated manually. **(C)** *Fcgr2b* gene score over pseudotime for individual trajectories and **(D)** flow cytometry analysis of FcγRIIB expression at indicated time points. **(E-I)** Congenically marked naïve CD8^+^ P14 cells were transferred into C57Bl/6 naïve recipient mice followed by LCMV Armstrong infection. FcγRIIB^-^ and FcγRIIB^+^ effector P14 cells were flow sorted from the spleen at 7 d p.i. and transferred into infection matched recipients. Transferred cells were isolated from the spleen and liver 30 d p.i. **(E)** Experimental schematics. **(F)** Total number of T_CM_, T_EM_ and T_RM_ P14 cells generated from FcγRIIB^-^ and FcγRIIB^+^ precursors and **(G)** Representative flow plots of CD69 and CD62L expression. **(H)** CD69 expression of FcγRIIB^-^ and FcγRIIB^+^ T cell progeny in the liver 30 d p.i. **(I)** Proportion of T_CM_ or T_EM_ and T_RM_ cells formed by FcγRIIB^-^ and FcγRIIB^+^ transferred cells. Data is pooled from 2 independent experiments with n=5 mice each. In **(D)** symbols represent mean. In **(F, H, I)** symbols represent individual mice. Bars represent mean. * p≤0.05, ** p≤ 0.01, *** p≤0.001, two-tailed Student’s t test.

Given the concomitant lack of *Fcgr2b* accessibility in T_MP-2_ and T_RM_ cells, we hypothesized that we may be able to identify T_RM_-poised precursors based on the surface expression of this marker. To observe FcγRIIB expression dynamics *in vivo*, we transferred P14 cells into LCMV infected mice and analysed FcγRIIB expression on P14 cells at various times p.i. Notably, the dynamics of *Fcgr2b* gene score in our predicted pseudotime trajectories closely reflected FcγRIIB expression by flow cytometry in the putative T_RM_, T_CM_, and T_EM_ cell populations over the course of infection (**Figure 3C and 3D**). Additionally, the comparison of FcγRIIB expression in memory T cell populations demonstrated increased expression in T_CIRC_ cells over time, with highest expression detected in the liver T_EM_ cell subset (**Figure S3B and S3C**).

We next asked whether the lack of FcγRIIB expression marked a population of precursor cells that might preferentially give rise to T_RM_ cells. For this, P14 cells were isolated from the spleen 7 d after LCMV infection and sort-purified populations of FcγRIIB^-^ or FcγRIIB^+^ P14 cells were transferred into infection-matched recipients (**Figure 3E**). At 30 d post-transfer, we observed a global reduction of FcγRIIB^+^ T cells (**Figure 3F**) consistent with their increased apoptotic potential (Morris et al., 2020), while FcγRIIB expression remained consistent as per the transferred population (**Figure S3D**). Interestingly, we found that the progeny of FcγRIIB^+^ cells displayed an increased proportion of T_CM_ and T_EM_ cells and an impaired conversion to the CD69_+_ T_RM_ cell phenotype (**Figure 3G-3I**). Accordingly, FcγRIIB^-^ showed an increased propensity to form T_RM_ cells at the detriment of either T_CIRC_ cell population in the liver. Together, these data indicate that differential FcγRIIB expression allows the identification of effector cells that appear to be differentially poised to navigate distinct memory T cell trajectories.

### T_RM_ cells share a core epigenetic signature across tissues

Our data thus far demonstrates that liver T_RM_ cells display a distinct epigenetic signature compared to their circulating counterparts derived from both the liver and spleen. Despite considerable variation of T_RM_ cells between organs, these cells exhibit a core transcriptional signature that is shared across tissues and species (Kumar et al., 2017; Mackay et al., 2016; Milner et al., 2017). Therefore, we sought to understand the extent of epigenetic similarity between CD8^+^ T_RM_ cells in different tissues and for this we compared liver T_RM_ cells to those derived from the skin. These populations represent extremes in T_RM_ cell-associated phenotypic and transcriptional variation, as driven by differential responsiveness to the cytokine TGF-β. Beyond upregulation of the integrin CD103, TGF-βgoverns a suite of transcriptional changes in the TGF-β-responsive skin T_RM_ cell population, in addition to restraining their functional capacity (Christo et al., 2021).

To define epigenetic features that are conserved between liver and skin T_RM_ cells, we utilized a model of HSV skin infection to induce a skin T_RM_ cell population. To this end, CD45.1^+^ gBT-I T cells specific for HSV gB_498-505_ were transferred into C57BL/6 mice that were infected with HSV on the skin flank. At 14 and 30 d post HSV infection, gBT-I T_RM_ cells were isolated from the skin as defined by the expression of CD69 and CD103, alongside circulating gBT-I T cells (T_CIRC_) derived from the skin-draining lymph node (dLN) (**Figure S4A**). Sort-purified cells were subjected to scATAC-seq and data was subsequently integrated with the aforementioned LCMV T cell dataset comprising spleen naïve and memory T cell subsets, and liver effector and memory T cell subsets (**Figure 4A**). As expected, liver and skin T_RM_ cells clustered separately, reflective of the major differences in their tissue microenvironment and phenotype. Differences were also observed in the clustering of T_CIRC_ cell populations, with clusters being defined by the model of infection (**Figure S4B**). Heatmap visualization of gene scores highlighted genes that were uniquely regulated in each cluster, as well as genes that were shared between skin (C2) and liver (C3) T_RM_ cells, including *Xcl1, Cdh1, Acvr1b,* and *Tnfsf11* (*Rankl*) (**Figure 4B**). As expected, skin T_RM_ cells had increased accessibility in genes related to TGF-β signalling (*Tgfbr2*, *Tgfbr3*), skin homing and adhesion molecules (*Ccr4*, *Ccr8*, *Itgae*) and genes previously shown to be preferentially expressed in skin T_RM_ cells (*Cish*, *Litaf*, *Pdcd1*, *Havcr2*). The loss in accessibility in genes that antagonize T_RM_ cell development (*Klf2, S1pr1)* could be observed within both T_RM_ cell clusters, together with an increased gene score in *Xcl1* and *Chn2*, genes identified as part of the core T_RM_ gene signature that is shared across organs (Mackay et al., 2016) (**Figure 4C and S4C-S4D**).

**Figure 4.**
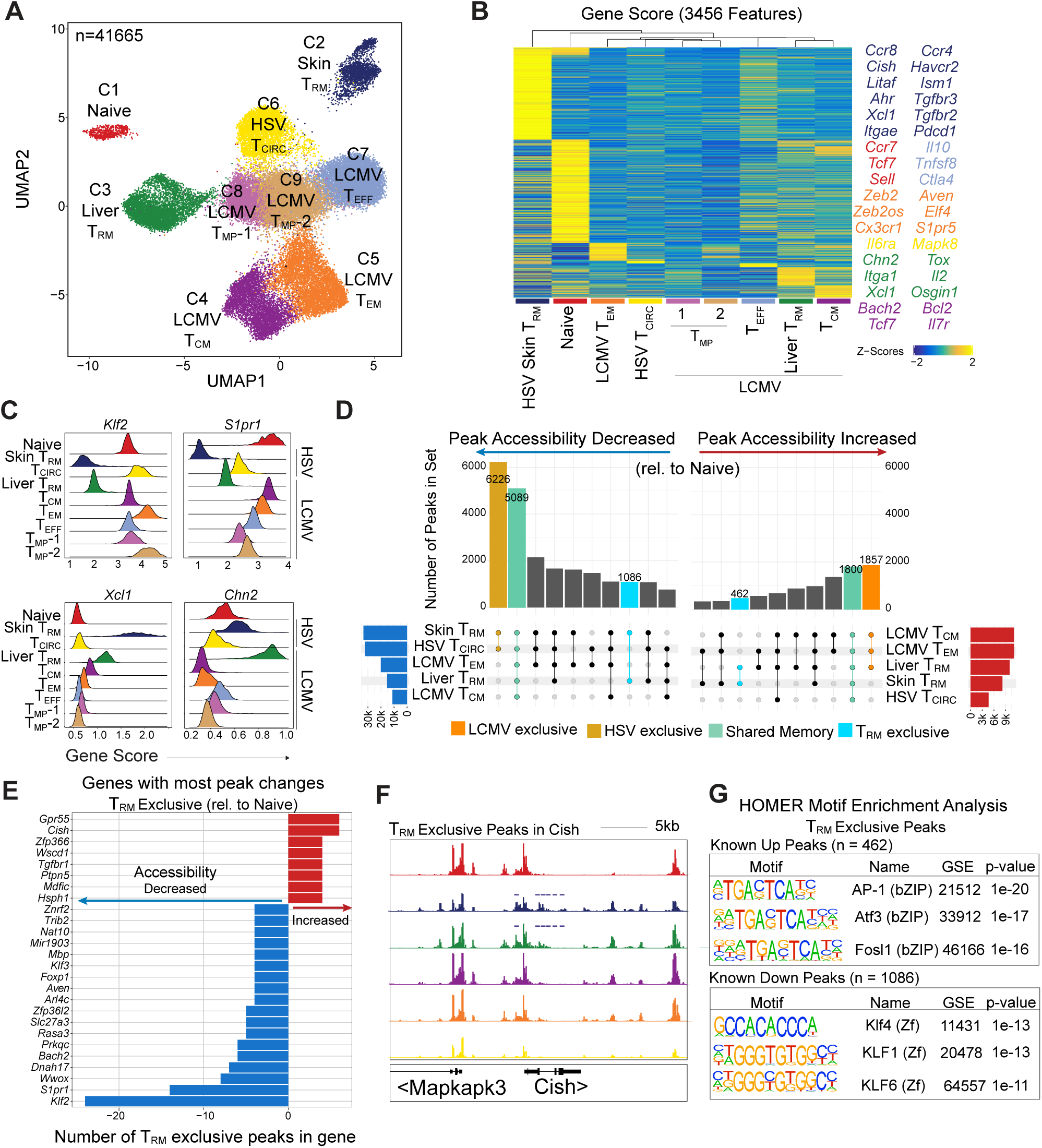
T_RM_ cells share a common epigenetic signature across tissues. **(A-G)** Congenically marked naïve CD8^+^ P14 cells were transferred into C57Bl/6 naïve recipient mice followed by LCMV Armstrong infection. P14 cells were flow sorted: from the liver at 7 d p.i., CD69- and CD69^+^ at 14 d p.i. and T_CM_, T_EM_ and T_RM_ from the spleen and liver at 30 d p.i. Congenically marked naïve CD8^+^ gBT-I cells were transferred into C57Bl/6 naïve recipient mice followed by HSV infection. gBT-I cells were flow sorted: from the axillary LN and from the skin (CD69^+^CD103^+^) at 14 and 30 d p.i., and scATACseq was performed. **(A)** UMAP projection and **(B)** Marker gene heatmap identifying genes that are uniquely accessible in each cluster; notable marker genes for each cluster annotated manually. **(C)** Histogram distribution of cluster gene scores for *Klf2, S1pr1, Xcl1* and *Cnh2.* **(D)** UpSet plot of shared peak sets in memory cells compared to naïve cells. **(E)** Common genes with most peak changes in T_RM_ cells relative to naïve cells. **(F)** Genome track of *Cish* in cluster aggregated scATAC-seq data (height normalized). **(G)** HOMER motif enrichment analysis of shared T_RM_ cluster peaks.

We next sought to determine if T_RM_ cells residing different organs had conserved accessibility in *cis*-regulatory elements. We identified significant changes in peaks relative to naïve T cells (C1) for all clusters and determined the extent to which sets of peaks were common across different clusters. Similar to the conservation of peaks in LCMV-induced memory subsets (**Figure 1D**), there was a strong conservation of *cis*-regulatory elements that exhibited changes in accessibility in all memory clusters relative to naïve (1,800 increased, 5,089 decreased) (**Figure 4D**). Across all peaks with significant changes in peak accessibility, 462 were increased and 1,082 decreased exclusively in both liver and skin T_RM_ cells. Mapping those 1,544 peaks back to genes revealed the genes with the most numerous changes in *cis*-regulatory element accessibility; expectedly, *Klf2* and *S1pr1* exhibited the most losses in peaks accessibility in T_RM_ cells. Among the genes with the most gains in *cis*-regulatory element accessibility were *Gpr55*, a G protein-coupled receptor that negatively regulates IEL T cell migration (Sumida et al., 2017) and *Cish* (**Figure 4E and 4F**), a negative regulator of TCR signalling (Palmer et al., 2015). Interestingly, we also observed conserved gains in peak accessibility at the *Tgfbr1* locus in both skin and liver T_RM_ cells (**Figure 4E**), which is intriguing given the opposing effect of TGFβ signalling on regulating T_RM_ cell development in these tissues (Christo et al., 2021).

To further understand gene-level accessibility changes unique to T_RM_ cells, we identified genes with significant changes in gene score relative to naïve T cells that did not appear in any other memory cluster (**Figure S4E**). In total, there were 64 genes with differential accessibility exclusively observed in both skin and liver T_RM_ cells. The 41 genes with increased accessibility in T_RM_ cells included the residency-associated chemokine *Xcl1*, cytokines *Il22* and *Tnfsf10,* and genes associated with cell adhesion (*Cd93*, *Gpr55*) or modulation of cytokine and TCR signalling (*Cish*, *Socs2*, *Tnfsf9*, *Rgs1*). We also analyzed whether T_RM_ cell-exclusive *cis*-regulatory elements shared transcription factor motifs that may control gene expression associated with changes in accessibility (**Figure 4G**). We observed a broad enrichment of motifs belonging to transcription factors from the bZIP family, with AP-1 motifs contributing to 30-40% of the 462 peaks with increased accessibility. In contrast, a reduction in KLF motifs was the most significantly enriched within the 1,062 peaks with decreased accessibility, in line with the role of KLF2 in antagonizing T_RM_ cell development (Skon et al., 2013). Together, these data indicate that a conserved epigenetic signature defines the T_RM_ cell population.

### Local microenvironment shapes the epigenome and promotes site-specific T_RM_ cell development

While T_RM_ cells shared an epigenetic signature across organs, it is well known that T_RM_ cells in different tissues exhibit discordant phenotypes and are regulated by distinct molecular cues (Christo et al., 2021; Fonseca et al., 2020; Frizzell et al., 2020; Kumar et al., 2017). Using our scATAC-seq data, we sought to determine the changes in chromatin landscape between T_RM_ cells from different organs and identify unique transcriptional regulators that may account for such differences. First, to determine the extent to which the chromatin state of skin and liver T_RM_ cells diverge, we compared with differential peak accessibility and gene scores between skin and liver T_RM_ cells (relative to T_CIRC_) (**Figure S5A**). In line with increased *P2rx7* mRNA levels (Mackay et al., 2016) (**Figure S5B**) and its known requirement for liver T_RM_ cell development (Stark et al., 2018), the *P2rx7* locus displayed increased accessibility in liver T_RM_ cells. In addition, we observed increased *Ahr* and *Ccr8* gene scores in skin T_RM_ cells, with a similar trend observed at the mRNA level (**Figure S5A and S5B**), fitting with the known roles for AHR and CCR8 in skin T_RM_ cell development (McCully et al., 2018; Zaid et al., 2014). Accordingly, molecular signature analysis of enriched motifs (MSigDB) in skin versus liver T_RM_ cells indicated participation of TGF-β signalling in skin T_RM_ cells, while liver T_RM_ cell motifs displayed enrichment in pathways related to IFN signalling (**Figure S5C**), confirming previous findings on the dependency of these respective cytokines for T_RM_ cell formation (Christo et al., 2021; Hirai et al., 2020; Holz et al., 2020; Mackay et al., 2013).

Next, we sought to identify putative T_RM_ cell regulators by combining transcriptional data from GSE70813 and epigenetic data to reveal transcription factors with both increased RNA expression and motif accessibility in a tissue-specific manner (**Figure 5A**). Compared to liver T_RM_ cells, skin T_RM_ cells had increased expression and accessibility in AP-1 family members, including JUN, JUNB, JUND, FOS, FOSB, FOSL1, and FOSL2, suggesting that one or more of these factors may specifically influence skin T_RM_ cell development. Of note, these comparisons also uncovered certain motifs with increased accessibility in skin compared to liver T_RM_ cells, but did not exhibit observable differential gene expression between these two T_RM_ subsets, such as BACH2. Accordingly, gene expression analysis comparing T_RM_ and T_CIRC_ cell subsets demonstrated that skin T_RM_ cells showed increased gene expression for *Fos, Fosb, Fosl1* and *Fosl2*, in addition to a small reduction in *Bach2* expression when compared to other T cell subsets, including liver T_RM_ cells (**Figure S5D**). This indicated a potential role for these transcription factors in skin T_RM_ cells that is not reflected in transcriptional data. As a confirmatory approach, we generated the list of peaks with increased accessibility exclusively in skin T_RM_ cells when compared to liver T_RM_ cells (relative to respective T_CIRC_ cell populations) and used the HOMER motif analysis to determine motif enrichment within skin T_RM_-exclusive peaks (**Figure 5B**). Here, AP-1 motifs were found in around 40% of peaks and the BACH2 motif was also amongst the top 10 enriched in the 2,663 peaks evaluated. When observed across memory T cell subsets, our data revealed that skin T_RM_ cells display the highest motif accessibility for FOS, FOSB, FOSL1, and BACH2 relative to all memory populations sequenced, indicating a putative role for these transcription factors in skin T_RM_ cell development (**Figure 5C and S5E-S5F**).

**Figure 5.**
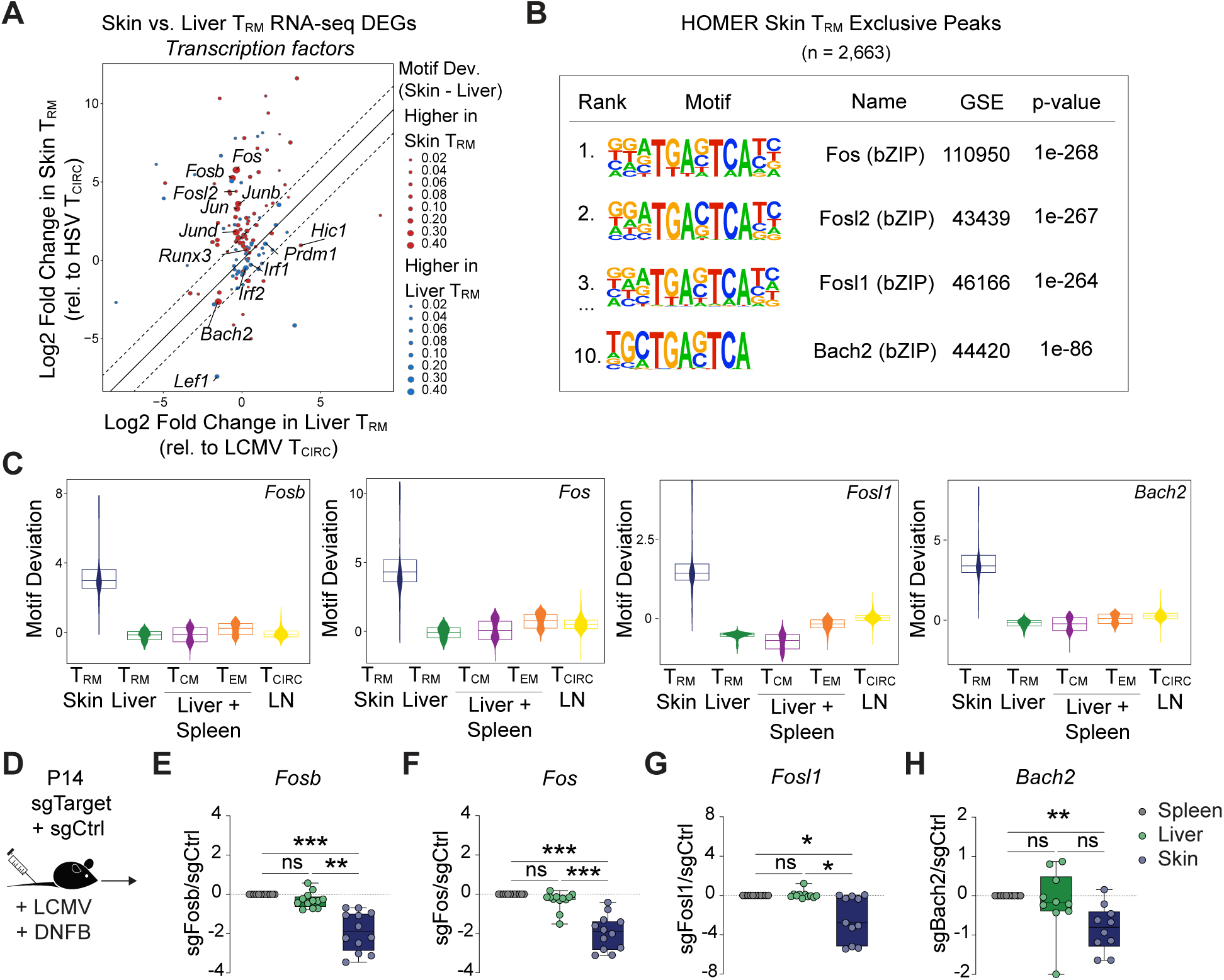
Tissue-specific epigenetic signatures depicts transcriptional regulators of T_RM_ cell development. **(A-C)** Congenically marked naïve CD8^+^ P14 cells were transferred into C57Bl/6 naïve recipient mice followed by LCMV Armstrong infection. P14 cells were flow sorted: from the liver at 7 d p.i., CD69- and CD69^+^ at 14 d p.i. and T_CM_, T_EM_ and T_RM_ from the spleen and liver at 30 d p.i. Congenically marked naïve CD8^+^ gBT-I cells were transferred into C57Bl/6 naïve recipient mice followed by HSV infection. gBT-I cells were flow sorted: from the axillary LN and from the skin (CD69^+^CD103^+^) at 14 and 30 d p.i., and scATACseq was performed. **(A)** Transcription factors enriched in significant T_RM_ cell motif deviations were selected and paired with DEGs between skin and liver T_RM_ cells normalized to gene expression in T_CM_ cells from GSE70813. **(B)** HOMER motif enrichment analysis of skin T_RM_ exclusive peaks. **(C)** FOSB, FOS, FOSL1 and BACH2 motif deviations in indicated populations of memory T cells. **(D-H)** Distinct congenically marked naïve CD8^+^ P14 cells were *in vitro* activated, and ablation of specific targets was performed using CRISPR-Cas9. Cells were then transferred into LCMV infected recipients that were treated with DNFB on the skin. Transferred cells were isolated from the spleen, liver and skin 30 d p.i. **(D)** Experimental schematics. Log2 FC of cells edited with **(E)** sgFosb, **(F)** sgFos, **(G)** sgFosl1 and **(H)** sgBach2 in the indicated tissues relative to sgCtrl normalized to the spleen. Data is pooled from **(D-H)** 2 independent experiments with n=5-6 mice each. In **(E-H)** symbols represent individual mice. Box plots show the median, interquartile range and minimum/maximum whiskers. * p≤ 0.05, ** p≤ 0.01, *** p≤0.001, ns p>0.05, One-way ANOVA with Bonferroni post-test.

Based on these data, we hypothesized that FOS family members, specifically FOS, FOSB and FOSL1, may uniquely regulate skin T_RM_ cells, in addition to their established role in controlling TCR-induced genes and T cell expansion (Roychoudhuri et al., 2016). To test this, we used CRISPR-Cas9 to ablate either *Fos, Fosb, Fosl1* or *Fosl2* in effector P14 cells and then cells were co-transferred together with cells edited with a control guide, into LCMV-infected mice (**Figure 5D**). To induce skin T_RM_ cells following LCMV infection, mice were treated with 2,4-dinitrofluorobenzene (DNFB) on the skin as previously described (Frizzell et al., 2020). Ablation of *Fosb* led to a general decrease in memory P14 cell formation in comparison to respective controls 30 d p.i. (**Figure S5G**). To observe location-specific defects in memory T cell formation, we compared the number of *Fosb*-deleted and control-edited cells in the skin and liver to normalized splenic cell numbers, and found a dominant defect in skin T_RM_ cell formation (**Figure 5E**). Similarly, the deletion of *Fos* and *Fosl1* (**Figure 5F and 5G**) revealed a loss of the skin T_RM_ cells population, whereas the deletion of *Fosl2* did not impact memory T cell formation (**Figure S5H**), consistent with the lack of changes in motif accessibility observed in T_RM_ cells.

To next investigate the role of the transcriptional repressor BACH2 in skin T_RM_ cell formation, we ablated this transcription factor in effector P14 cells via CRISPR-Cas9 and co-transferred edited cells and control cells into LCMV-infected DNFB-treated recipient mice (**Figure 5H**). Akin to our findings above, we observed a pronounced defect in BACH2-deleted T cells specifically in the skin, as compared to those isolated from the spleen or liver (**Figure 5H and S5I**). Together, this demonstrates the utility of integrating transcriptional and epigenetic analysis to identify major regulators of tissue-specific T cell development. Further, scATAC-seq enabled the identification of differential BACH2 activity where RNA-seq could not, highlighting the utility of scATAC-seq in nominating novel transcriptional regulators of cell state.

### T_RM_ and T_EX_ cells are epigenetically distinct

A canonical feature of T_RM_ cells is their elevated expression of inhibitory receptors as compared to circulating memory T cells. This is particularly striking for skin T_RM_ cells, which share several phenotypic characteristics with T_EX_ cells generated in response to chronic viral infection, presenting similar reduced capacity for cytokine production (Christo et al., 2021). To directly compare T_EX_ and memory T cell subsets, we transferred congenically marked naïve CD8^+^ P14 cells into C57Bl/6 recipient mice infected with LCMV Armstrong or LCMV Clone-13 (Cl-13) infection, a widely used model for inducing CD8^+^ T cell exhaustion, or gBT-I cells after HSV infection (**Figure 6A**). Based on exhaustion and memory T cell markers (**Figure S6A**), UMAP visualization revealed four major clusters (**Figure 6A**), with skin gBT-I T_RM_ cells exhibiting similar expression of the checkpoint molecules PD1 and TIM3 as splenic P14 cells isolated from mice infected with LCMV Clone-13 (Cl-13) (**Figure 6B**). In contrast, liver T_RM_ cells generated in response to acute LCMV do not express PD1 or TIM3 to the same extent **(Figure 6A and 6B)**. Further, tumor-infiltrating lymphocytes (TIL) have also been reported to have a T_RM_ cell-like transcriptional profile (Djenidi et al., 2015; Malik et al., 2017; Milner et al., 2017; Nizard et al., 2017; Park et al., 2019; Savas et al., 2018). The seemingly convergent phenotypic profiles of T_RM_ and T_EX_ cells has led to speculation that T_EX_ and T_RM_ cell lineages are related (Blank et al., 2019). It is unclear however, the extent to which the chromatin state of T_RM_ and T_EX_ cells overlap, or when these subsets diverge during T cell differentiation.

**Figure 6.**
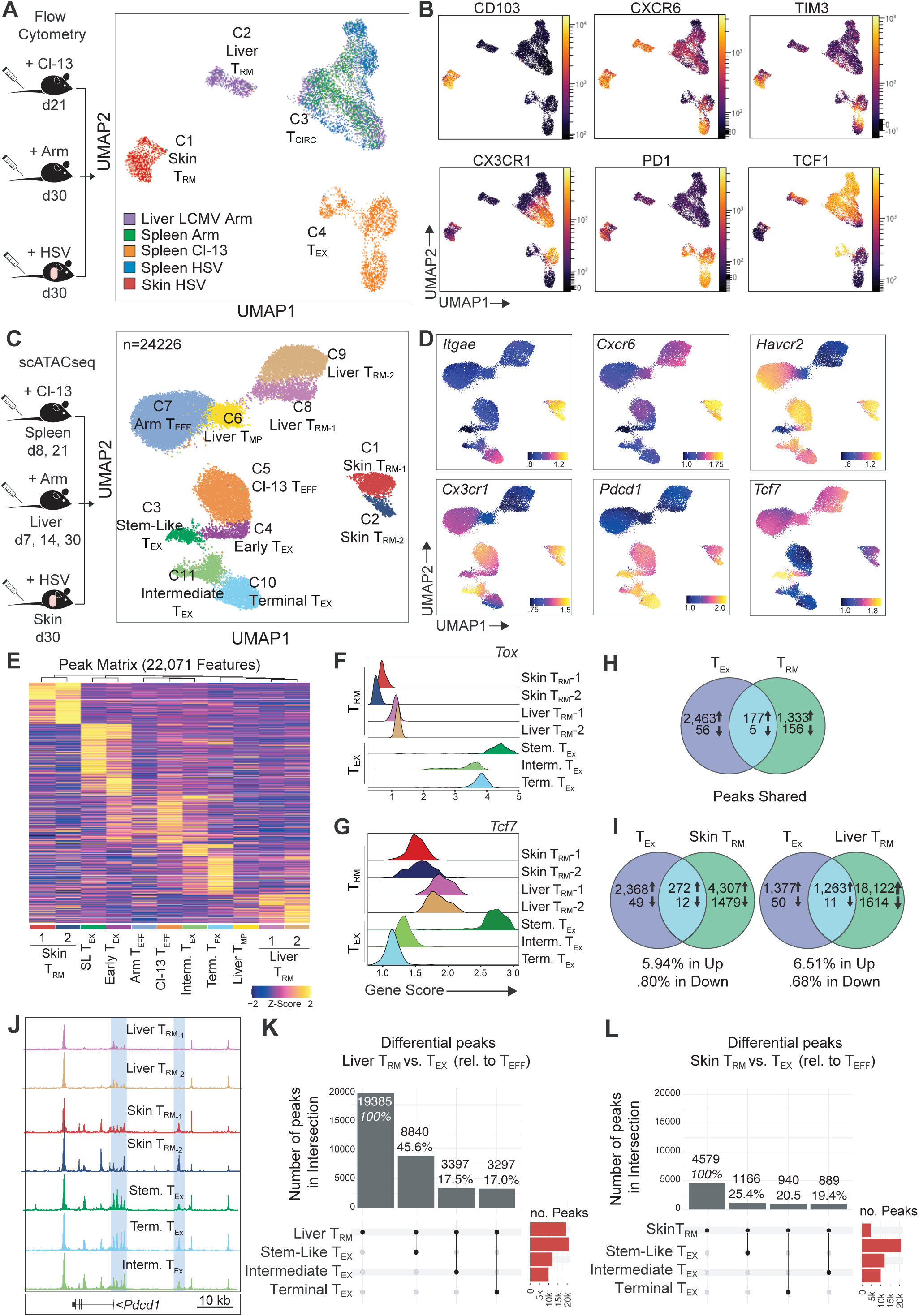
T_RM_ cells are epigenetically distinct from T_EX_ cells. **(A, B)** Congenically marked naïve CD8^+^ P14 cells were transferred into C57Bl/6 recipient mice followed by LCMV Armstrong or LCMV Clone-13 infection. Congenically marked naïve gBT-I cells were transferred into C57Bl/6 recipient mice followed by HSV infection. Spleen, liver and skin of infected mice were harvested at indicated timepoints for flow cytometry. **(A)** Experimental schematics and UMAP projection of the indicated T cell populations in tissues based on flow cytometric analysis. **(B)** UMAP depicting expression of indicated markers across clusters. **(C-L)** Congenically marked naïve CD8^+^ P14 or gBT-I cells were transferred into C57Bl/6 recipient mice followed by LCMV Armstrong or HSV infection respectively. A separate group of naïve C57Bl/6 mice were infected with LCMV Clone-13. Total P14 cells were flow sorted from the liver at 7 d p.i., CD69^+^ at 14 d p.i. and T_RM_ cells at 30 d p.i. gBT-I cells were flow sorted from the skin (CD69^+^CD103^+^) at 30 d p.i. Endogenous gp33^+^ cells were flow sorted from the spleen of Clone-13 infected mice at 8 and 21 d p.i. scATACseq was performed in isolated populations. **(C)** Experimental schematic and UMAP projection based on scATACseq analysis. **(D)** UMAP depicting relative gene accessibility (gene score) across clusters. **(E)** Heatmap identifying peaks that are uniquely accessible in each cluster relative to all clusters. **(F)** *Tox* and **(G)** *Tcf7* gene scores in T_RM_ and T_EX_ clusters. **(H)** Venn diagram depicting similar peaks with increased or decreased accessibility in T_RM_ and T_EX_ cells and **(I)** specific peaks shared with liver or skin T_RM_ and T_EX_ cells. **(J)** *Pdcd1* genome tracks of cluster aggregated scATAC-seq data (height normalized); peaks with qualitative height differences highlighted. **(K)** UpSet plot of skin or **(L)** liver T_RM_ cluster shared peak sets with exhausted T cell subsets compared to effector T cells.

To assess epigenetic differences between these subsets, we analyzed scATAC-seq data from gp33 tetramer^+^ CD8^+^ T cells from LCMV Cl-13 infected mice at 8 and 21 d p.i. from GSE188670 (Daniel et al., 2021) together with our scATAC-seq data of liver and skin T_RM_ cells, as generated by LCMV Armstrong and HSV infections, respectively (**Figure 6C**). These integrated datasets revealed that T_EX_ cells separated into 5 distinct clusters; cells isolated at 8 d p.i. were primarily classified as C4 or C5, and cells isolated at 21 d p.i. mostly inhabited C3, C10, and C11 (**Figure S6B**). To separate stem-like T_EX_ cells, intermediate T_EX_ cells, and terminal T_EX_ cells (Raju et al., 2021), we used *Havcr2*, *Cx3cr1*, *Pdcd1,* and *Tcf7* gene scores, identifying C3, C11 and C10 respectively (**Figure 6D**). Notably, LCMV Cl-13-induced T_EX_ cells clustered separately from cells isolated from LCMV Armstrong infected hosts at all stages of differentiation, highlighting distinct chromatin states for T_RM_ and T_EX_ cells (**Figure 6E**).

Next, we sought to compare gene and motif accessibility for transcription factors commonly associated with T cell exhaustion in T_EX_ and T_RM_ cells. We found that *Tox* gene scores in both skin and liver T_RM_ cells were reduced in comparison to all T_EX_ subsets (**Figure 6F**), supporting low *Tox* expression in T_RM_ cells (**Figure S6C**). Additionally, similar to the expression at the protein level, increased *Tcf7* gene score was observed in stem-like T_EX_ cells and liver T_RM_ cells (**Figure 6G**), consistent with the increased stemness observed in these populations that maintain increased differentiation capacity. Motif deviation for several T_EX_-associated transcription factors, TCF1, IRF4, and EOMES, were highest in T_EX_ clusters (**Figure S6D**). RUNX3, however, exhibited the highest motif deviation in the skin T_RM_ cell population, consistent with its required role for residency in several tissues (Milner and Goldrath, 2018) (**Figure S6D**).

To more broadly understand the shared epigenetic regulation of T_RM_ and T_EX_ cells, we compared peaks with significantly increased or decreased accessibility in each T cell subset relative to T_EFF_ cells generated after LCMV Armstrong infection. First, we compared shared T_RM_ peaks (significant peaks present in both liver and skin T_RM_) to shared T_EX_ cell peaks (significant peaks present in all three T_EX_ subsets isolated 21 d p.i.), i.e. T_RM_ and T_EX_ programs, respectively. This analysis showed that among the 1,510 peaks that have increased accessibility in both T_RM_ subsets, only 177 were shared with the T_EX_ program (2,640 peaks) (**Figure 6H**). We assigned these 177 peaks to the nearest gene and ordered the genes with the most assigned peaks; *Slc24a5*, a cation exchanger, and *Sesn3*, a stress-sensing protein that can promote NKR recognition in CD8^+^ T cells (Pereira et al., 2020), were the most commonly assigned genes with 3 peaks each (**Figure S6E**). Comparing the T_EX_ program to individual skin or liver T_RM_ cell peaks sets yielded similar results (272 shared of 4,579 total peaks in skin T_RM_ cells; 1,263 shared of 19,385 total peaks in liver T_RM_ cells) with skin T_RM_ and liver T_RM_ cells sharing 5.94% and 6.51% peaks with T_EX_ cells, respectively (**Figure 6I**). Interestingly, 2 of the 272 peaks that skin T_RM_ cells share with T_EX_ are nearest to the *Pdcd1* gene; one peak ∼23kb from the TSS of *Pdcd1* is a known T_EX_ enhancer that mediates sustained PD1 expression, previously thought to be specific to T_EX_ cells (Pauken et al., 2016; Sen et al., 2016) (**Figure 6J**). The presence of this peak in both skin T_RM_ and gp33-specific CD8^+^ T_EX_ cells could potentially explain the constitutive expression of PD1 in both T cell subsets.

Finally, we then compared the peak set of skin and liver T_RM_ clusters with each individual T_EX_ cluster to determine which T_EX_ cell population most closely shared the epigenetic features of T_RM_ cells. Interestingly, skin T_RM_ cells exhibited similar amounts of significant increased or decreased peaks with the three T_EX_ subsets, while liver T_RM_ cells shared mostly peaks with stem-like T_EX_ cells (**Figure 6K and L**), potentially due to stem-like T_EX_ cells being the most “memory-like” of the T_EX_ cell subsets. Altogether, our results indicate that even though T_RM_ cells may share phenotypic similarities with exhausted T cells, epigenetic analyses demonstrate major differences in gene accessibility changes throughout development that define specific memory or exhausted T cell subsets.

## Discussion

Here, we utilized scATAC-seq to examine epigenetic changes that occur over the course of the T cell response against acute, local and systemic, or chronic viral infections. Our data defines the epigenetic variation between individual T cell subsets at various stages of infection and reveals an early divergence of memory precursors destined for a circulating or tissue-resident cell fate. We demonstrate that T_RM_ cells are an epigenetically distinct T cell subset that share a conserved epigenetic signature across organs, as well as tissue-specific epigenetic variation. Together, our findings highlight the dramatic changes in chromatin landscape that underlie cellular differentiation, as well as the resolution of scATAC-seq to finely distinguish individual populations within the CD8_+_ T cell pool (Lareau et al., 2019; Satpathy et al., 2019). Differences in chromatin accessibility across effector and memory T cells can be further interrogated in our public genome browser, allowing visualization of scATAC-seq reads in specific gene loci to investigate T cell biology.

Our data supports a model by which memory T cell fate is determined early post infection, supporting previous findings that demonstrate early fate decision for T_RM_ cell generation in the skin and gut (Kok et al., 2020; Milner et al., 2020). Using trajectory analyses, we modelled gene and transcription factor motif accessibility over time in effector, memory precursors and memory T cells, providing a genome-wide view of changes in the epigenome over the course of memory T cell differentiation. Importantly, by looking at genes with increased accessibility in the T_RM_-poised memory precursor cluster we identified major epigenetic divergences defining early effector commitment to the T_RM_ and T_CIRC_ cell populations. Specifically, we uncovered the differential expression of FcγRIIB between T_RM_ and T_CIRC_ cells that is retained from their respective precursors. These findings allowed the selection of an effector population with enhanced capacity for generating each of those subsets, adding to the previous characterization of the role of FcγRIIB in triggering apoptosis to limit T cell mediated immunity (Morris et al., 2020).

We identified a common epigenetic signature conserved between T_RM_ cells from different organs, consisting of key gene regulatory networks that contribute to T cell retention, in addition to reduced ability to traffic in the blood and secondary lymphoid organs. In addition to this conserved T_RM_ cell program (Kumar et al., 2017; Mackay et al., 2016; Milner et al., 2017), cells residing in different organs exhibit divergent phenotypes and functional capacities as shaped by extrinsic cues in their distinct tissue microenvironments (Fonseca et al., 2020; Frizzell et al., 2020). We leveraged our chromatin accessibility data to identify potential transcriptional regulators that support tissue-specific T_RM_ cell formation and validated the role of several transcription factors including HIC1, as well as AP-1 factors, FOSB, BACH2, FOS, and FOSL1, adding to their previously established role in T cell memory by regulating the availability of AP-1 motifs to limit the expression of TCR-driven genes during T cell effector responses (Roychoudhuri et al., 2016; Yukawa et al., 2020). Together, these results support the concept that epigenetic differences can underlie tissue-specific modulation of gene expression and that interrogation of these changes can reveal novel, tissue-specific transcriptional regulation.

T_RM_ cells generated in response to acute infection and T_EX_ cells in tumors and in chronic infection share considerable phenotypic overlap (Christo et al., 2021; Milner et al., 2017, 2020). Skin T_RM_ cells, in particular, display restricted functional capacity and increased inhibitory receptor expression, characteristics that are typically associated with the T_EX_ cell lineage (Christo et al., 2021; Park et al., 2019). A longstanding question has been whether the phenotypic and functional overlap between skin T_RM_ cells and T_EX_ cells is indicative of a shared epigenetic state or a convergence of cell types. Here, epigenetic analysis of T_RM_ and T_EX_ subsets demonstrated that despite expression similarities in the expression of certain co-inhibitory receptors such as PD-1, the epigenetic state of T_RM_, T_CIRC_ and T_EX_ subsets are ultimately distinct. Our detailed analysis demonstrated that despite similarities in the accessibility of certain genes, the majority of the epigenetic changes occurring in T_EX_ cells are not present in skin or liver T_RM_ cells.

Understanding the regulation and differentiation of T_RM_ cells is critical to informing the design of therapies that aim to modulate tissue immunity. Our data demonstrates that T_RM_ cells are an epigenetically distinct T cell subset that arise from epigenetically poised precursors generated early after infection, as well as revealing novel regulators of T_RM_ cell differentiation. Altogether, our results provide critical insights into T_RM_ cell differentiation and phenotype and will act as resource for further investigation into events that precede T_RM_, T_CIRC_ and T_EX_ cell differentiation, and epigenetic regulators that contribute to T_RM_ cell maintenance and function.

## Acknowledgements

We thank the Flow Cytometry Unit and Bioresources Facility at Peter Doherty Institute (University of Melbourne) for technical assistance and the Stanford Functional Genomics Facility for sequencing support. This work was supported by an Institutional Training Grant 5T32AI007290 (F.A.B.), a Howard Hughes Medical Institute and Bill & Melinda Gates International Research Scholarship OPP1175796 (L.K.M.), National Health and Medical Research Council (NHMRC) AP1113293 (L.K.M.), the National Institutes of Health (NIH) K08CA230188, U01CA260852, and UM1HG012076 (A.T.S.), the Parker Institute for Cancer Immunotherapy (A.T.S.), a Career Award for Medical Scientists from the Burroughs Wellcome Fund (A.T.S), and a Pew-Stewart Scholars for Cancer Research Award (A.T.S). L.K.M is a Senior Medical Research Fellow supported by the Sylvia and Charles Viertel Charitable Foundation.

## Author contributions

F.A.B., R.F, J.A.B., M.E., A.O., Y.Q., B.D. and K.E.Y. performed experiments and analyzed data; A.T.S. and L.K.M. provided supervision; F.A.B, R.F., A.T.S, and L.K.M. contributed to experimental design. F.A.B, R.F., A.T.S, and L.K.M. prepared the manuscript. A.T.S. and L.K.M. provided funding and led the research program.

## Supplementary Figure legends

**Figure S1 (Related to Figure 1).**
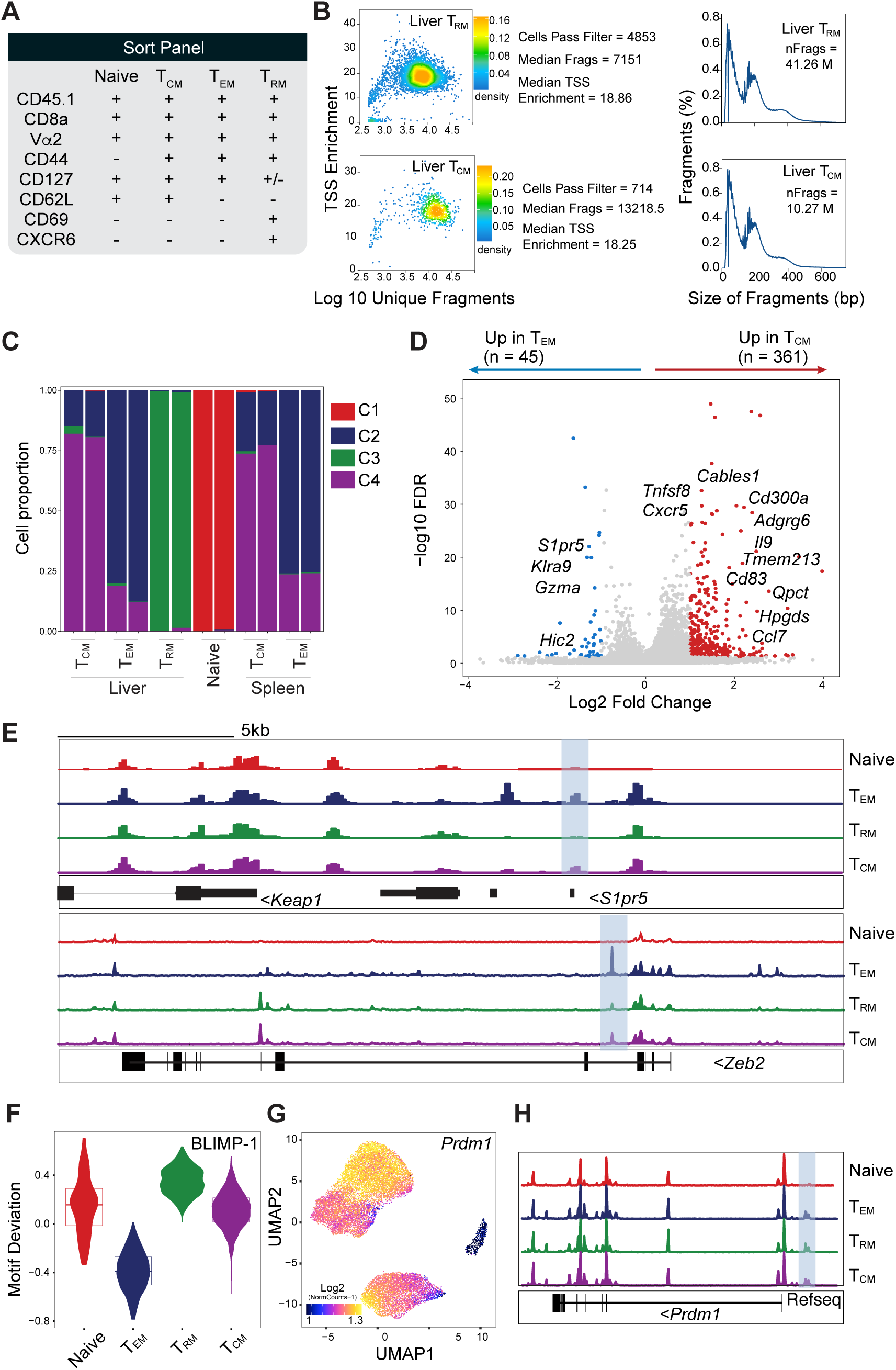
Epigenetic variation of memory T cell subsets following LCMV infection. **(A-H)** Congenically marked naïve CD8^+^ P14 cells were transferred into naïve recipient mice followed by LCMV Armstrong infection. T_CM_ (CD62L^+^ CD69^-^), T_EM_ (CD62L^-^ CD69^-^) and T_RM_ (CD62L^-^CD69^+^) cells were flow sorted from the spleen and liver 30 d p.i. and scATACseq was performed. **(A)** Summary of antigen expression among CD8^+^ T cell subsets for each marker contained in the cell sort antibody panel. **(B)** Representative quality control plots. **(C)** Cluster composition by sample identity based on CD8^+^ T cell subsets sorted. **(D)** Gene score volcano plots identifying genes with different accessibility (log2 FC > 1, FDR > 10) between T_EM_ and T_CM_ memory clusters; notable genes annotated manually. **(E)** Genome tracks of *S1pr5 and Zeb2* (height normalized). **(F)** BLIMP-1 motif deviation, **(G)** UMAP depicting relative gene accessibility (gene score) across clusters and **(H)** cluster aggregated genome track of scATAC-seq data (height normalized).

**Figure S2 (Related to Figure 2).**
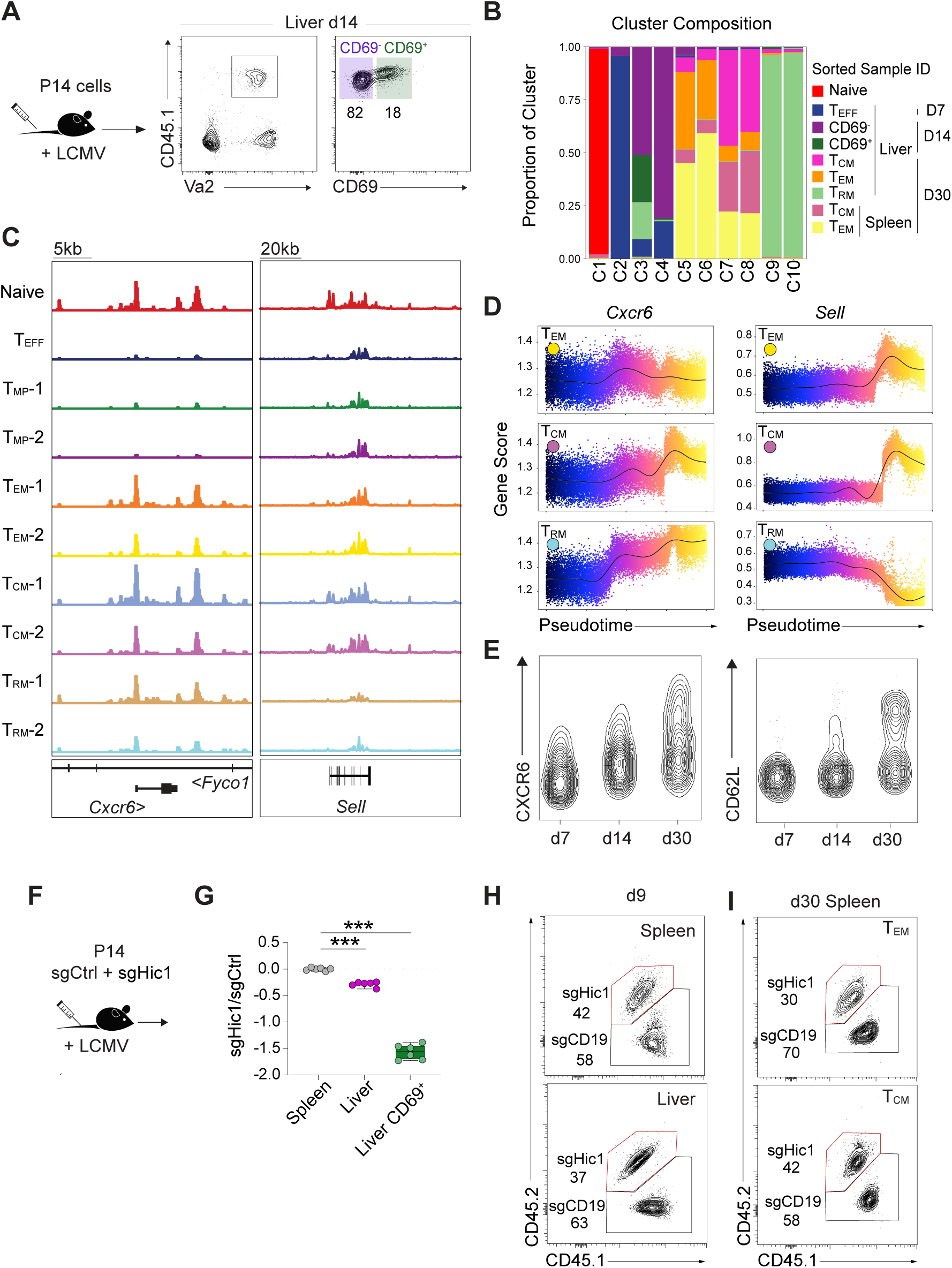
Epigenetic and phenotypic variations drive development of distinct memory T cell subsets. **(A-D)** Congenically marked naïve CD8^+^ P14 cells were transferred into naïve recipient mice followed by LCMV Armstrong infection. P14 cells were flow sorted: from the liver at 7 d p.i., CD69- and CD69^+^ at 14 d p.i. and T_CM_, T_EM_ and T_RM_ from the spleen and liver at 30 d p.i. and scATACseq was performed. **(A)** Experimental schematics and representative flow plots of sorted populations from the liver 14 d p.i. **(B)** Cluster composition by sample identity. **(C)** Genome tracks of *Cxcr6* and *Sell* (height normalized). **(D)** *Cxcr6* and *Sell* gene score over pseudotime for individual trajectories. **(E)** Flow cytometry analysis of CXCR6 and CD62L expression by CD8^+^ P14 cells after LCMV Armstrong infection at indicated time points. **(F-I)** Control (sgCtrl) or *Hic1* (sgHic1) ablation was performed using CRISPR-Cas9 in distinct congenically marked naïve CD8^+^ P14 cells. Cells were then transferred into LCMV infected recipients and isolated from the spleen and liver 9 and 30 d p.i. **(F)** Experimental schematics and **(G)** Log2 FC of sgHic1 and sgCtrl indicated cell subsets normalized to the spleen at 9 d p.i. **(H-I)** Representative flow plots of transferred cells **(H)** in the spleen and liver 9 d p.i. and **(I)** in the indicated subsets in the spleen 30 d p.i. Data is representative of **(E)** 2 independent experiments with n=5 mice each and **(F-I)** 2 independent experiments with n=6-10 mice each. In **(G)** symbols represent individual mice. Box plots show the median, interquartile range and minimum/maximum whiskers. *** p≤0.001, One-way ANOVA with Bonferroni post-test.

**Figure S3 (Related to Figure 3).**
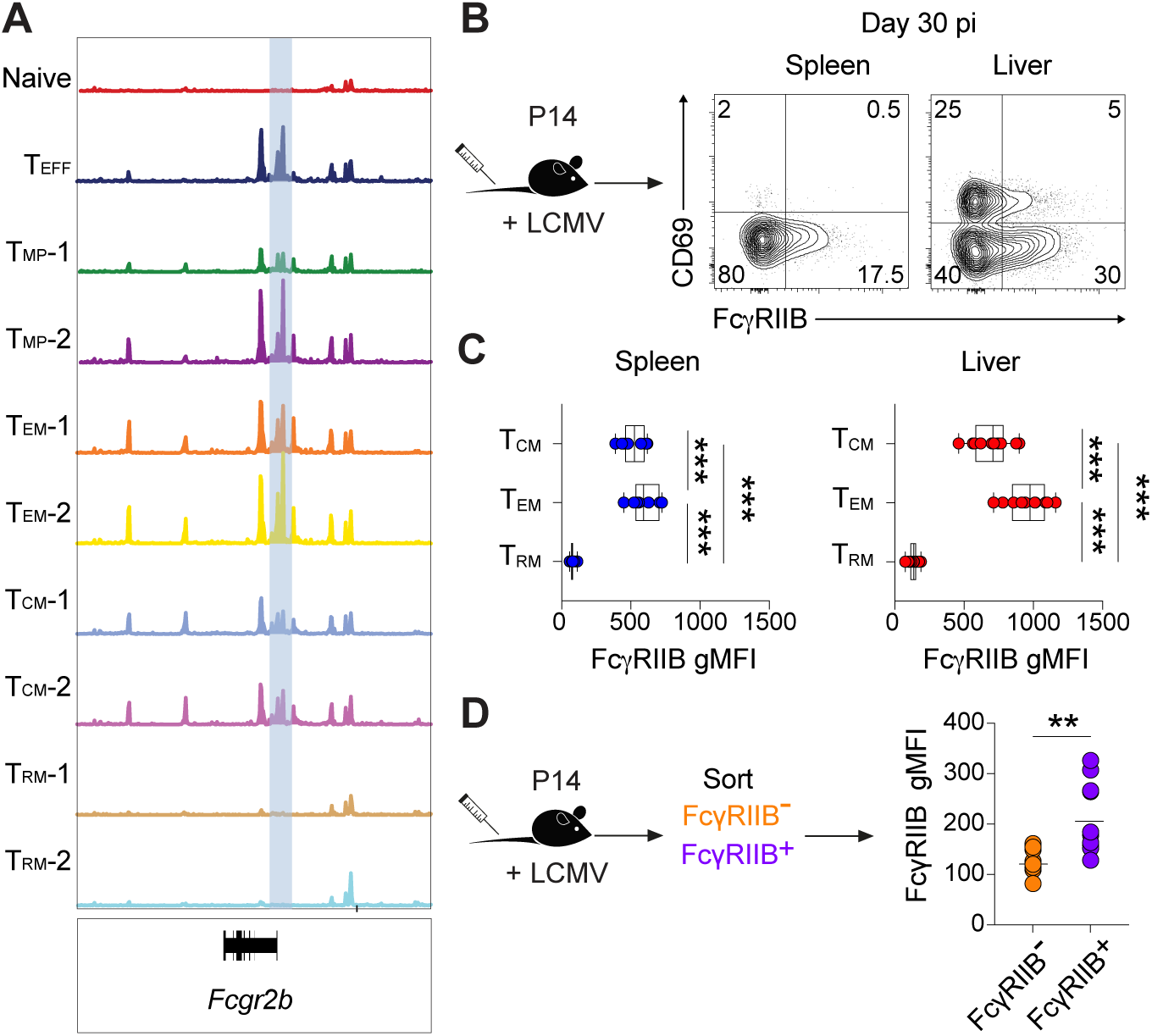
FcγRIIB expression reflects gene accessibility in T cell subsets. **(A)** Congenically marked naïve CD8^+^ P14 cells were transferred into naïve recipient mice followed by LCMV Armstrong infection. P14 cells were flow sorted: from the liver at 7 d p.i., CD69^-^ and CD69^+^ at 14 d p.i. and T_CM_, T_EM_ and T_RM_ from the spleen and liver at 30 d p.i. and scATACseq was performed. Genome tracks of *Fcgr2b* (height normalized). **(B, C)** Congenically marked naïve CD8^+^ P14 cells were transferred into naïve recipient mice followed by LCMV Armstrong infection and isolated 30 days later from the spleen and liver. **(B)** Experimental schematics and representative flow plots of CD69 and FcγRIIB expression in the spleen and liver. **(C)** FcγRIIB expression by T_CM_, T_EM_ and T_RM_ subsets. **(D)** Congenically marked naïve CD8^+^ P14 cells were transferred into naïve recipient mice followed by LCMV Armstrong infection. FcγRIIB^-^ and FcγRIIB^+^ effector P14 cells were flow sorted from the spleen at 7 d p.i. and transferred into infection matched recipients. Transferred cells were isolated from the liver 30 d p.i. Experimental schematics and FcγRIIB expression in isolated progeny. Data is pooled from 2 independent experiments with n=5 mice each. In **(C, D)** symbols represent individual mice. Bars represent mean. ** p≤0.01, *** p≤0.001, One-way ANOVA with Bonferroni post-test.

**Figure S4 (Related to Figure 4).**
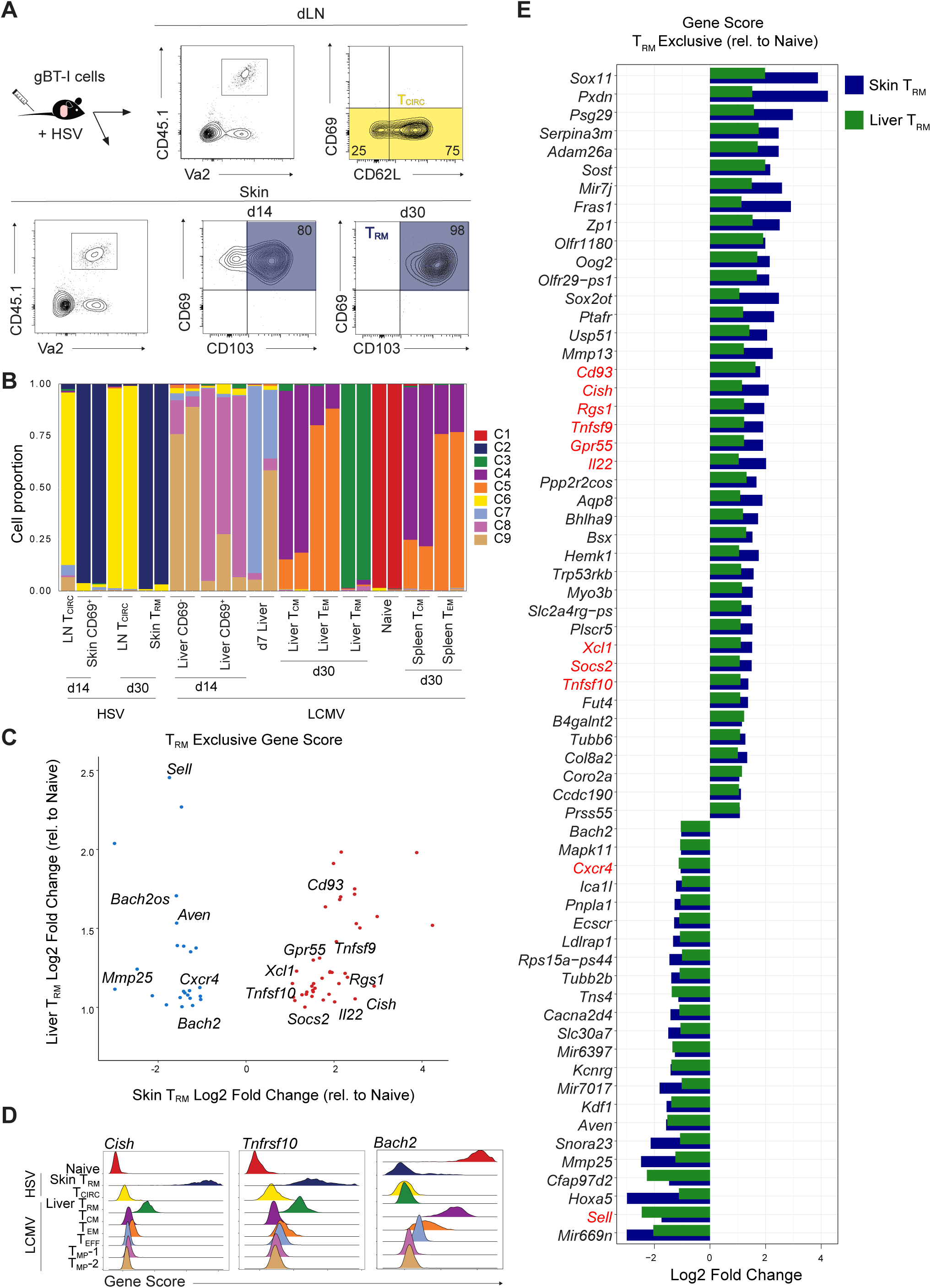
T_RM_ cells display a conserved epigenetic profile across tissues. **(A-E)** Congenically marked naïve CD8^+^ P14 T cells were transferred into naïve recipient mice followed by LCMV Armstrong infection. P14 T cells were flow sorted: from the liver at 7 d p.i., CD69- and CD69^+^ at 14 d p.i. and T_CM_, T_EM_ and T_RM_ from the spleen and liver at 30 d p.i. Congenically marked naïve CD8^+^ gBT-I T cells were transferred into naïve recipient mice followed by HSV infection. gBT-I T cells were flow sorted: from the skin-draining (axillary) LN and from the skin (CD69^+^CD103^+^) at 14 and 30 d p.i., and scATACseq was performed. **(A)** Experimental schematic and representative flow plots of sorted populations from skin and skin-draining LN 14 and 30 d p.i. **(B)** Cluster composition by sample identity. **(C)** Dot plot depicting liver and skin T_RM_ exclusive gene scores relative to naïve cells. **(D)** *Cish, Tnfrsf10 and Bach2* gene scores in individual indiciated clusters. **(E)** Genes with significant gene score differences exclusively in skin and liver T_RM_ cells relative to naïve.

**Figure S5 (Related to Figure 5).**
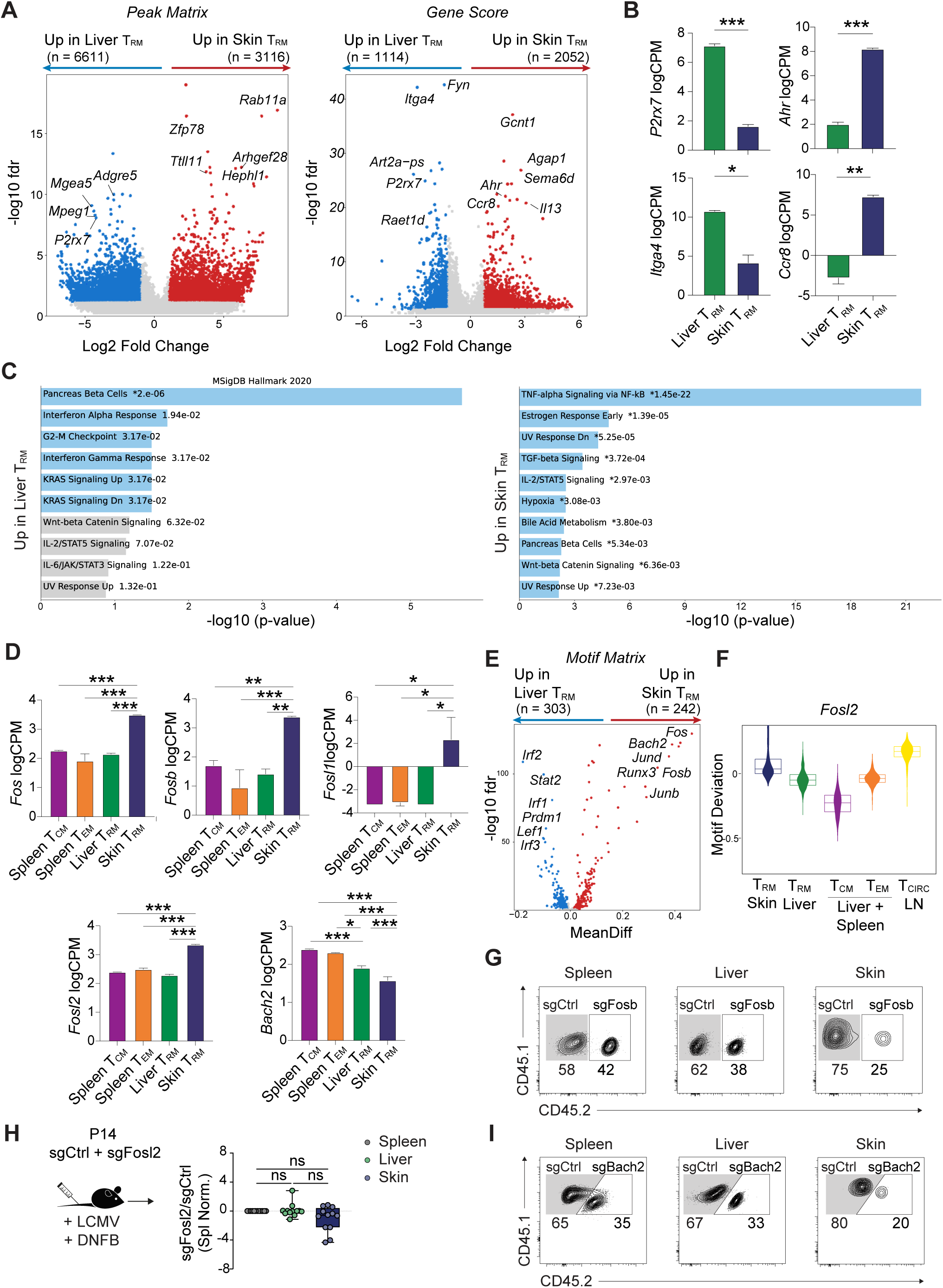
Epigenetic profile reveals tissue-exclusive pathways and requirements for T_RM_ cell development. **(A, C, E, F)** Congenically marked naïve CD8^+^ P14 T cells or gBT-I T cells were transferred into naïve recipient mice followed by LCMV Armstrong or HSV infection, respectively. P14 T_RM_ cells were flow sorted from the liver and gBT-I T_RM_ cells were flow sorted from the skin at 30 d p.i. scATACseq was performed in isolated populations. Volcano plot depicting differences between skin and liver T_RM_ cells exclusive gene peaks and gene scores. **(A)** Volcano plot depicting differences between skin and liver T_RM_ cells exclusive peaks and gene scores. **(B)** Indicated genes log counts per million (logCPM) in skin and liver T_RM_ cells from GSE70813. **(C)** Pathway analysis of liver and skin T_RM_ cell exclusive motifs enriched in the 2020 Molecular Signature Database (MSigDB). **(D)** Indicated genes log counts per million (logCPM) in spleen T_CM_ and T_EM_ cells, liver T_RM_ cells and skin T_RM_ cells from GSE70813. **(E)** Volcano plot depicting skin and liver T_RM_ cell exclusive motifs. **(F)** FOSL2 motif deviation in indicated populations of memory T cells. **(G-I)** Distinct congenically marked naïve CD8^+^ P14 cells were *in vitro* activated, and ablation of specific targets was performed using CRISPR-Cas9. Cells were then transferred into LCMV infected recipients that were treated with DNFB on the skin. Transferred cells were isolated from the spleen, liver and skin 30 d p.i. **(G)** Representative flow plots of sgFosb and sgCtrl transferred cells. **(H)** Log2 FC of sgFosl2 and sgCtrl cells. **(I)** Representative flow plots of sgBach2 and sgCtrl transferred cells. Data is pooled from **(G-I)** 2 independent experiments with n=5-6 mice each. In **(G-I)** symbols represent individual mice. Box plots show the median, interquartile range and minimum/maximum whiskers. * p≤ 0.05,** p≤ 0.01, *** p≤0.001, One-way ANOVA with Bonferroni post-test.

**Figure S6 (Related to Figure 6).**
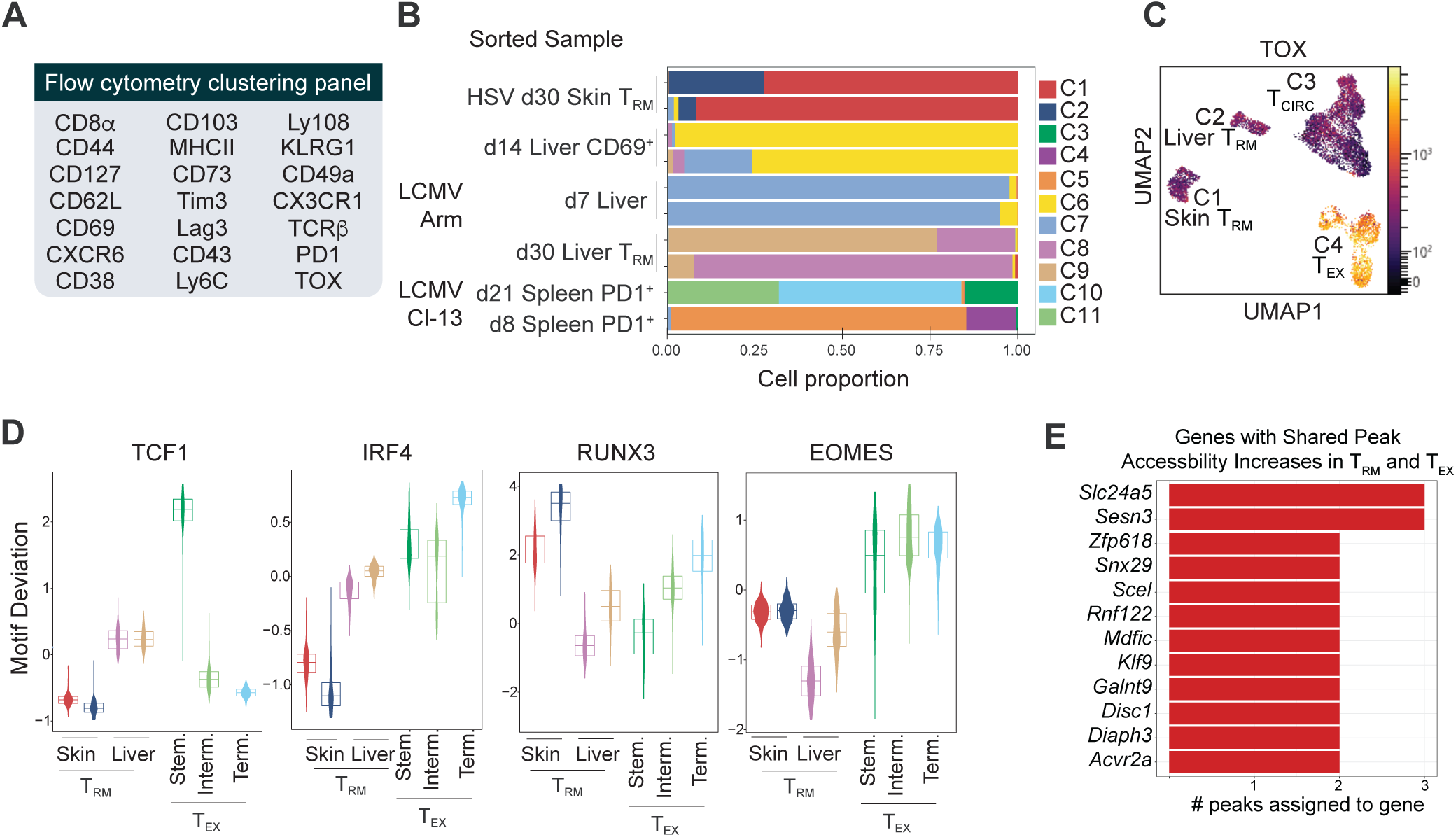
T_RM_ and T_EX_ cell subsets are epigenetically distinct. **(A, C)** Congenically marked naïve CD8^+^ P14 cells were transferred into recipient mice followed by LCMV Armstrong or LCMV Clone-13 infection. Congenically marked naïve gBT-I cells were transferred into recipient mice followed by HSV infection. Spleen, liver and skin of infected mice were harvested at indicated timepoints for flow cytometry. **(A)** Summary of the antibody panel utilized for clustering CD8^+^ T cell subsets in Fig. 6a. **(B, D, E)** Congenically marked naïve CD8^+^ P14 cells were transferred into recipient mice followed by LCMV Armstrong. Congenically marked naïve gBT-I cells were transferred into recipient mice followed by HSV infection. Naïve mice were infected with LCMV Clone-13. Endogenous gp33^+^ cells were flow sorted from the spleen at 8 and 21 d p.i. scATACseq was performed in isolated populations. **(B)** Cluster composition by sample identity. **(C)** Shown is TOX expression in T cell clusters. **(D)** *Tcf7, Irf4, Runx3* and *Eomes* motif deviation in indicated T cell clusters. **(E)** Gene assignment of peaks with increased accessibility in both T_RM_ and T_EX_ cell subsets relative to Arm T_EFF_ cells.

## Material and Methods

### Mice

C57BL/6, gBT-I.CD45.1. P14.CD45.1 and P14.CD45.1.2 mice were bred in the Department of Microbiology and Immunology at the University of Melbourne. Six-to eight-week-old C56BL/6 were used for experiments. All experiments were approved by the University of Melbourne Animal Ethics Committee.

### Adoptive cell transfers, infections and DNFB treatment

For naïve transgenic T cell transfers, cells were isolated from lymph nodes and spleen and transferred intravenously (i.v.) to C57BL/6 mice at 5×10^4^ cells per recipient. Skin infections were done by skin scarification with 1×10^6^ plaque-forming units (PFU) of HSV-1 KOS as described^81^. LCMV Armstrong infections were done by intraperitoneal injection of 2×10^5^ PFU to establish acute infections. LCMV Clone-13 experiments were done by i.v. injection of 1×10^6^ PFU to establish chronic infections. For treatment with 1-Fluoro-2,4-dinitrobenzene (DNFB), mice were shaved and depilated before treatment with 15 μl of DNFB (Sigma-Aldrich) diluted at 0.25% in acetone:oil (4:1) on the skin as described(Mackay et al., 2012b) or on the ears 3 d after LCMV infection.

### Organ processing, flow cytometry, and cell sorting

Spleens were processed through metal meshes into single-cell suspensions followed by red blood cell lysis. Skin samples were excised and incubated at 37°C for 90 min in dispase (2.5 mg/ml; Roche) or in liberase (0.25 mg/ml; Sigma) followed by separation of epidermis and dermis. Chopped samples were incubated at 37°C for 30 min in collagenase III (3 mg/ml; Worthington). Liver samples were excised and meshed into single-cell suspensions through 70 μm meshes. Leukocytes were isolated using Percoll (35%; Sigma Aldrich). Single cell suspensions were stained with conjugated antibodies for flow cytometry or cell sorting. For intracellular staining of cytokines and transcription factors, cells were fixed and permeabilized using the Foxp3 Transcription factor Staining buffer set (Invitrogen) as per manufacturer’s instructions. The following antibodies from BD Biosciences, Biolegend, Cell signalling or Thermo Fisher Scientific were used: anti-CD45.1 (A20), anti-CD45.2 (104), anti-CD8α (53-6.7), anti-CD8β (YTS1 56.7.7), anti-CD3 (500A2), anti-Vα2 (B20.1), anti-CD44 (IM7), anti-CD127 (A7R34), anti-CXCR6 (SA051D1), anti-CX3CR1 (SA011F11), anti-CD62L (MEL-14), anti-CD69 (H1.2F3), anti-CD103 (2E7), anti-PD-1 (29F.1A12), anti-Ly6C (HK1.4), anti-TCRβ (H57-597), anti-TIM-3 (RMT3-23), anti-CD43 (1B11), anti-Ly108 (330-AJ), anti-CD38 (90), anti-CD49a (HMa1), anti-CD32b (AT130-2), anti-gp33 tetramer, anti-TOX (TXRX10), anti-TCF7 (C63D9), anti-MHC-II (M5/114.15.2), anti-CD73 (TY/11.8), anti-LAG3 (C9B7W), anti-KLRG1 (2F1). Cell viability was determined using Ghost Dye Red 780 (Tonbo Biosciences). Flow cytometry was performed on a LSRFortessa (BD Biosciences) or an Aurora (Cytek) and analyzed with FlowJo software (TreeStar) or Omiq. For cell sorting experiments, P14 and gBT-I cells were isolated from the spleen, liver and skin as indicated and sorted using a FACSAria III (BD Biosciences).

### CRISPR-Cas9 Gene Editing of CD8^+^ T cells

Single guide RNA (sgRNA) targeting: *CD19* (5’-AAUGUCUCAGACCAUAUGGG-3’), *Hic1 (*5’-AGUGUGCGGAAAGCGCGGAG-3’, 5’-CUUGUGCGACGUGAUCAUCG-3’*), Fos (*5’-TGTCACCGTGGGGATAAAGTTGG-3’, 5’-GGTCTGCGATGGGGCCACGGAGG-3’*), Fosb (*5’-AGACAGGTACTGAGACTCGGCGG-3’, 5’-GTTGACCCTTATGACATGCCAGG-3’*), Fosl1 (*5’-GGAACCGGGACCGAGCTCCGGGG-3’, 5’-GCTGCGCGGGGCGACCGTACGGG-3’*), Fosl2 (*5’-GACGAGGTGTCAAAGTTCCCGGG-3’, 5’-GGACATGGAGGTGATCACTGTGG-3’*), Bach2 (*5’-TGCGCAGGAACTCAGCACAGCGG-3’, 5’-GATGTTGGCACAGTGGACTGTGG-3’) were purchased from Synthego (CRISPRevolution sgRNA EZ Kit). sgRNA/Cas9 RNPs were formed by incubating 0.3nmol of sgRNA with 0.6 ml Alt-R S.p. Cas9 nuclease V3 (10 mg/ml; Integrated DNA Technologies) for 10 min at room temperature. P14 cells were *in vitro* activated with anti-CD3 and anti-CD28 (5 μg/ml) for 24 hours. *In vitro* activated or naïve P14 cells were resuspended in 20 μl of P3 (P3 Primary Cell 4D-Nucleofector X Kit; Lonza), mixed with sgRNA/Cas9 RNP and electroporated using a Lonza 4D-Nucleofector system (CM137) as previously described (Nüssing et al., 2020). Cells were expanded for 72 hours in the presence of IL-2 (25U/ml; Peprotech). Naïve and *in vitro* activated edited cells were mixed at a 1:1 ratio and 5×10^5^ cells were transferred i.v. into LCMV-infected recipients.

### scATAC-seq library preparation and sequencing

Sorted T cell populations were thawed, then subjected to the 10x Chromium scATAC protocol (https://support.10xgenomics.com/single-cell-atac). In short, nuclei were isolated and partitioned into gel-bead emulsions that allow barcoded transposition to happen at single cell scale. Following transposition, the emulsions were broken, the product was cleaned and libraries were prepared for Illumina sequencing. Libraries were sequenced on the Illumina HiSeq 4000.

### scATAC-seq computational analysis

Fastq files were trimmed, aligned to the mm10 reference genome, and deduplicated using the 10X genomics cellranger-atac count pipeline. Fragments files for each sample, containing the unique aligned reads passing filter for each cell barcode, were then loaded into ArchR for downstream analysis (Granja et al., 2021). Doublet identification and removal, cell calling, clustering, peak calling, and motif analysis was performed using the default ArchR workflow. After initial clustering, there was often a small cluster of contaminating non-T cells -- these were manually identified, removed, and the remaining cells were reclustered following the same procedure. Marker features were identified using Archr ‘getMarkerFeatures’. GeneScore visualizations were performed using the ArchR implementation of Magic imputation. Motifs enriched in specific peak sets were analyzed using the HOMER (Heinz et al., 2010) ‘findMotifsGenome.pl’ utility. For these analyses, the indicated peak set was compared to a background peak set consisting of the ArchR union peak set for that group of samples. Therefore, the motifs identified represent motifs enriched relative to the rest of the peaks in these samples, rather than motifs enriched in, for example, T cells in general.

### RNA-Seq Analysis

For TF expression analysis, previously generated RNA-seq count matrixes were analyzed using DESeq2 (Love et al., 2014) using the default parameters. T_RM_ samples were compared to their respective T_CIRC_ populations using the ‘results’ function with α =0.05.

### Statistical analysis

Statistical analyses were performed by one- or two-way analysis of variance (ANOVA) test followed by Bonferroni’s post-test or by two-tailed Student’s *t* test using Prism 9 (GraphPad) as indicated in figure legends. *P* values were represented by * *p* < 0.05; ** *p* < 0.01; *** *p* < 0.001; **** *p* < 0.0001; ns (not significant) *p > 0.05*. Results represent means ± SEM.

### Data availability

All original data is available from the corresponding author upon reasonable request. Sequencing data is available in the Gene Expression Omnibus database under accession code GSE199799. Source Data are provided in the online version of the manuscript. Code available upon request.

